# Synonymous codon substitutions regulate transcription and translation of an upstream gene

**DOI:** 10.1101/2022.08.05.502938

**Authors:** Anabel Rodriguez, Gabriel S. Wright, Taylor J. Lundgren, McKenze J. Moss, Jun Li, Tijana Milenkovic, Paul W. Huber, Matthew M. Champion, Scott J. Emrich, Patricia L. Clark

## Abstract

Synonymous codons were originally viewed as interchangeable with no phenotypic consequences. However, over the years a substantial body of evidence has demonstrated that some synonymous substitutions can perturb a variety of gene expression and protein homeostasis mechanisms, including translational efficiency, translational fidelity and co-translational folding of the encoded protein. To date, synonymous codon-derived perturbations have largely focused on effects within a single gene. Here we show that synonymous codon substitutions made far within an *E. coli* plasmid-encoded protein coding sequence frequently led to significant upregulation of a neighboring, upstream gene. Notably, in four out of nine synonymously recoded sequences, significant upregulation of the upstream gene arose due to cryptic transcription of the anti-sense strand. Surprisingly, cryptic transcription of the upstream gene readily bypassed its native transcriptional repression mechanism. Even more surprisingly, translation of this upstream gene correlates closely with the subset of its mRNA transcribed from the cryptic internal promoter, rather than its total mRNA level. These results suggest that synonymous codons in bacteria may be under selection to both preserve the amino acid sequence of the encoded gene while also avoiding internal sequence elements that significantly perturb transcriptional and translational regulation of neighboring genes.

## INTRODUCTION

The degeneracy of the genetic code means that each protein sequence can be encoded by an astronomically large number of distinct DNA coding sequences. On average, each amino acid is encoded by three synonymous codons, meaning that a single protein sequence of 200 amino acids can be encoded by ∼3^200^ (∼10^94^) unique sequences. Because synonymous substitutions do not alter the encoded amino acid sequence, synonymous codons were historically assumed to be interchangeable, with no effect on cell function. However, it has long been known that some synonymous codon substitutions can indeed impact gene expression and/or protein homeostasis (1, 2), for example by causing significant changes in transcription level (3), translation initiation efficiency (4), translational elongation rate (5), amino acid misincorporation frequency (6) and co-translational protein folding mechanisms (7–9). Some synonymous mutations reduce cell fitness (10, 11) and/or increase disease susceptibility (12–14).

The growing number of mechanisms sensitive to synonymous codon substitutions has led to an explosion of synonymous codon usage measures. These measures are used to compare codon usage of one gene to another, variations in codon usage within a single coding sequence, and likewise to test for correlations between codon usage and other phenomena, including gene expression level and protein folding efficiency (12). Many codon usage measures build upon the foundational observation that highly expressed *E. coli* genes tend to be enriched in common codons and depleted in rare codons, which led to the development of the codon adaption index (CAI) (1). Other, more recently developed codon usage measures include tAI (15, 16), nTE (17) and %MinMax (18, 19), which calculate codon usage relative to tRNA gene copy number, tRNA supply versus codon demand, and relative codon usage frequencies for a given amino acid sequence, respectively. However, there have been few direct comparisons of the abilities of these measures to predict outcomes such as gene expression level or protein folding efficiency (see (7), however, for an exception related to co-translational folding).

To date, most studies of codon usage have focused on the effects of synonymous codon substitutions within a single gene. These effects span not just translation but also transcription. For example, in eukaryotes, synonymous codons substitutions can alter chromatin structure, which can impact transcription level (20). The entire genome of *E. coli* and other bacteria is densely packed, with many genes immediately adjacent to, or even overlapping, each other (21, 22). The proximity of genes in *E. coli* raises the possibility that synonymous substitutions in one gene could significantly affect transcription and/or translation of a neighboring gene.

Here we used an established *E. coli* plasmid expression system based on the *E. coli* tetA/tetR operon to test the effects of synonymous mRNA sequence recoding on gene expression. We designed and expressed a variety of synonymous sequences encoding the antibiotic resistance protein chloramphenicol acetyltransferase (CAT) under the control of a commonly used *tet* ON/*tet* OFF overlapping divergent promoter system (23). Surprisingly, we found that synonymous recoding of *cat* mRNA readily introduced sequences within *cat* that led to enhanced cryptic transcription of the divergent, upstream *tetR* coding sequence. This cryptic transcription of *tetR* originates from a start site within *cat*, far upstream from the canonical *tetR* transcription start site, and bypasses the native *tetR* repression mechanism. Even more surprisingly, TetR protein production correlated closely with the level of *tetR* cryptic mRNA but was uncorrelated with total *tetR* mRNA level. The discovery of significant intergenic regulation of transcription and translation originating within the coding sequence of a neighboring gene expands the range of phenotypic effects sensitive to synonymous codon usage, identifying a novel mechanism by which synonymous codons can more broadly alter gene expression.

## MATERIAL AND METHODS

### Design of *cat* synonymous sequences

To complement an analysis of the wild type and previously described synonymous Shuf1 *cat* coding sequences (8), we designed, synthesized, and analyzed eight additional synonymous *cat* sequences (**Supplementary Methods**). Six of these sequences (MM-Harm, HPMM-Harm, CAI-Harm, tAI-Harm, nTE-Harm, and AllModel) were designed to correlate closely (“harmonize”) with a WT cat codon usage profile; these were created using the CHARMING codon harmonizing algorithm (**Table S1**) (24). Except for AllModel, each of these harmonized sequences was designed such that its codon usage profile correlates closely with the WT profile according to a single codon usage measure but is weakly correlated with the WT profile as calculated using other codon usage measures (**Table S2)**. In contrast, the AllModel sequence was designed to cross-correlate with multiple codon usage profiles. The GA sequence was generated using the online GeneOptimizer tool (GeneArt; www.thermofisher.com), selecting *E. coli* as the host organism (25). In contrast to all other *cat* constructs tested here, the overall codon composition of WTscr is identical to WT CAT; it was designed by randomly swapping synonymous codon positions in the wild type coding sequence. All *cat* coding sequences tested here preserve the wild type CAT protein sequence.

### Construction of *cat* mRNA synonymous sequences

The designed *cat* synonymous sequences were cloned into plasmid pKT, under the control of a titratable *tet* promoter (23). All oligonucleotides (**Supplementary Tables S3, S4**) were ordered from Integrated DNA Technologies (IDT). MEGAWHOP (26) and FastCloning (27) were used to clone synonymous *cat* sequences into the pKT plasmid. Initially, these sequences each included a C-terminal ssrA degradation tag; this tag was subsequently removed using FastCloning to generate the final constructs used here. To prevent CAT translation, the *cat* ribosome binding site was mutated from GAAGGA to TTCTCT using FastCloning. To test whether some *cat* coding sequences introduce new TetR binding sites, the *cat* expression plasmid was modified using FastCloning to insert codons 40-220 from four *cat* mutants (WT, GA, CAI, and WTscr) 287 nucleotides downstream of the *cat* stop codon. The sequence of each construct used here was confirmed by Sanger sequencing.

### CAT and TetR protein quantification

Saturated overnight cultures of *E. coli* KA12 were prepared by inoculating 5 mL of Luria Broth (LB) containing 100 ug/mL ampicillin with a single colony followed by growth overnight with shaking at 37°C. The resulting saturated culture was used to inoculate to 3 mL of LB containing 100 μg/mL ampicillin and 600 ng/mL tetracycline (tc) to an OD_600_ of 0.05. Cultures were grown with shaking at 37°C for 3 h, then cells were pelleted, resuspended to OD_600_ 10 in SDS loading dye, boiled for 20 min, spun for 1 min at 15,000 rpm, and 15 µL of cleared cell lysate (or resuspended insoluble material) was separated by electrophoresis on a 12% polyacrylamide gel. Protein bands were transferred to a nitrocellulose membrane and probed with primary antibodies [mouse anti-His (Qiagen) to detect CAT; mouse anti-TetR (MoBiTec); mouse anti-GAPDH (Invitrogen)]. The membrane was washed three times, then incubated with goat anti-mouse Alexa647-labeled secondary antibody (Invitrogen) followed by measurement of fluorescence emission (excitation: 628 nm, emission: 676 nm) across the blot. CAT and TetR protein bands were quantified using ImageJ, corrected to the loading control GAPDH, and normalized to WT to compare bands across multiple blots.

### Quantification of *cat* and *tetR* mRNA

mRNA abundance was determined using reverse transcriptase quantitative polymerase chain reaction (RT-qPCR). Saturated overnight cultures of *E. coli* KA12 containing 100 μg/mL ampicillin were used to inoculate 12-well plates containing 3 mL of LB, 100 μg/mL ampicillin, and 600 ng/mL tetracycline (tc). Wells were inoculated to OD_600_ 0.05 and grown using a Synergy H1 microplate reader (BioTek) with continuous double-orbital shaking at 37°C, for 3 hours. An aliquot of cells was diluted to OD_600_ 0.3 and 500 µL of diluted cells was added to 1 mL of RNAprotect Bacteria Reagent provided in the RNeasy Mini Kit (Qiagen; catalog #74104). Protocols provided in the “RNAprotect Bacteria Reagent Handbook 2015” were used for RNA extraction (“Protocol 1”), on-column DNase digestion (“Appendix B”), and RNA purification (“Protocol 7”). RNA concentration was determined by absorbance at 260 nm on a NanoVue spectrophotometer. To remove any contaminating genomic or plasmid DNA, 10 µL of RNA sample (1.0-1.9 µg) were digested with 2 units of RNase-free DNase (Ambion/Invitrogen) in 10 µL of RNase-free DNase buffer at 37°C for 30 min. Samples were spiked with 1 µL of DNase and incubated at 37°C for another 30 min. To denature the DNase, 1 µL of 100 mM EDTA was added and samples were incubated at 65°C for 10 min.

Reverse transcription (RT) was performed with the iScript RT kit (BioRad) using 10 µL (0.50-0.95 µg) of RNA digested with DNase. Total RT reaction volume was 20 µL (10 µL RNA, 4 µL 5x buffer, 2 µL random hexamer primers, 1 µL reverse transcriptase). A control lacking reverse transcriptase was included for all mutants. The RT reaction was performed on a MyCycler (BioRad) using the following parameters: 5 min at 25°C, 30 min at 42°C, 5 min at 85°C.

qPCR reaction volume per sample was 5 µL total (1 µL of five-fold diluted cDNA, 60 nM primers (IDT), and 2.5 μL SsoAdvanced SYBR Green Supermix (BioRad)). Using a liquid handling robot, 5 µL was dispensed in technical triplicate for each CAT mutant into a hard-shell, 96-well, thin-wall PCR plate (BioRad). Amplification was performed using a CFX96 Touch Real-Time PCR Detection System (BioRad). Primers used for amplification were: *cat* (Forward primer: GCATCACCATCACCATCACCATAAC; Reverse primer: GATAAAACTCAAAATGCTCACGGCG), total tetR (Forward primer: CGTAAACTCGCCCAGAAGCTA; Reverse primer: ATCTTGCCAGCTTTCCCCTTC), cryptic tetR (Forward: TAAGCAGCTCTAATGCGCTGT; Reverse: TTAATTTCGCGGGATCGGCTA), and GAPDH reference gene (Forward primer: CGGTACCGTTGAAGTGAAAGAC; Reverse primer: ACCAGTTGCTTCAGCGAC). (and GAPDH reference gene (Forward primer: CGGTACCGTTGAAGTGAAAGAC; Reverse primer: ACCAGTTGCTTCAGCGAC). PCR efficiency for each primer set: *cat* (103.98%; R^2^ = 0.996), total *tetR* (103.58%; R^2^ = 0.987), cryptic *tetR* (98.88%; R^2^ = 0.989), *GAPDH* (95.60%, R^2^ = 0.998).

To calculate fold change, ΔCt values were calculated for each biological replicate per mutant (ΔCt = Ct of target - Ct of GAPDH). To calculate ΔΔCt for all constructs, the ΔCt of WT was subtracted from the ΔCt for each biological replicate (ΔΔCt = ΔCt target - ΔCt WT) to minimize variation in the data due to plate-to-plate variation between biological replicates. This results in a ΔΔCt value for each biological replicate per construct. The fold change is calculated as 2^-ΔΔCt^ per biological replicate per construct. Fold change was averaged across three biological replicates; mean fold change and standard deviation are plotted relative to WT.

### Plasmid DNA quantification

*E. coli* KA12 transformed with a pKT-*cat* expression plasmid was grown as described above for 3 hours without inducer. A 1 mL culture aliquot was pelleted at 4,600x*g* for 10 min at 4°C. The supernatant was discarded, and cell pellets were resuspended in PBS pH 7.4 (1.47 mM KH_2_PO_4_, 7.81 mM Na_2_HPO_4_, 137 mM NaCl, 2.68 mM KCl). Cells were diluted 1:100 in PBS and 100 µL were lysed as described in (28) qPCR reactions were carried out as described above for mRNA quantification using the same CAT and GAPDH primers and 1 μL of lysate in place of cDNA. Mean-fold change was calculated as described above except ΔΔCT was calculated by subtracting the average ΔCT WT (ΔΔCT = ΔCT target – average ΔCT WT). All qPCR reactions were performed on the same 96-well plate.

## RESULTS

### Design strategy for *cat* synonymous codon substitutions

We previously identified Shuf1, a synonymous recoding of the *E. coli cat* coding sequence that subtly alters the CAT protein native structure (8). As a next step towards sampling the large number of synonymous *cat* coding sequences and their effects on gene expression and/or protein homeostasis, we designed five additional synonymous *cat* coding sequences that each have low identity to the wild type coding sequence (**Table 1**) (and to each other; **Supplementary Tables S5, S6**) but correlate closely with (e.g., is “harmonized” to (24)) the profile for a codon usage measure of the wild type *cat* coding sequence (i.e. harmonized sequence) (**Figure 1A; Supplementary Table S2**). These harmonized sequences also have very similar %GC and total number of codon and nucleotide changes (**Supplementary Tables S5, S6**). The five codon usage measures we selected (CAI, tAI, nTE, %MinMax and High-phi %MinMax) are distinct (24) (see **Supplementary Methods**) but each is commonly used as proxy for translation elongation rate (12), which can affect translational efficiency and hence intracellular protein accumulation. We also created a sixth harmonized sequence designed to correlate as closely as possible to all five codon usage measures (AllModel; AllM). In addition to these six harmonized sequences, we also created a synonymous non-harmonized sequence using a more traditional codon optimization strategy (GA)(25), and another that scrambles the order of the wild type *cat* synonymous codons while retaining the wild type amino acid sequence (WTscramble; WTscr) (**Figure 1B**). As with our previous Shuf1 coding sequence, each of these eight total new recoded *cat* coding sequences retains the first 46 codons from wild type *cat* (**Figure 1A, B**) to minimize known effects of 5’ synonymous codon changes on translation initiation (4, 29–32). The complete sequence of each *cat* construct used in this study is listed in **Supplementary Methods**.

**Table 1.** Synonymous *cat* mutant sequence characteristics.

| CAT mutant | Spearman<br>Correlation<br>to WT | Changes relative to WT |  |  |  | %GC | CAI<sub>g</sub> |
| --- | --- | --- | --- | --- | --- | --- | --- |
|  |  | % codon | # codon | % nt | # nt |  |  |
| WT-CAT | - | 0 | 0 | 0 | 0 | 36 | 0.65 |
| %MinMax (**MM**) | 0.813 | 42.5 | 94 | 17 | 113 | 42 | 0.66 |
| High-Phi MinMax (**HPMM**) | 0.792 | 46.6 | 103 | 18.4 | 122 | 41 | 0.60 |
| normalized Translational<br>Efficiency (**nTE**) | 0.966 | 42.5 | 94 | 16.9 | 112 | 42 | 0.63 |
| tRNA Adaptation Index (**tAI**) | 0.944 | 50.2 | 111 | 20.2 | 134 | 40 | 0.60 |
| CAI (**CAI**) | 0.899 | 49.8 | 110 | 19.3 | 128 | 42 | 0.59 |
| AllModel (**AIIM**) | 0.832† | 44.3 | 98 | 17.2 | 114 | 40 | 0.60 |
| GeneArt (**GA**) | - | 38.9 | 86 | 16.4 | 109 | 41 | 0.85 |
| WTscramble (**WTscr**) | - | 49.8 | 110 | 20.5 | 136 | 36 | 0.65 |
| Shuf1 | - | 46.2 | 102 | 18.6 | 123 | 42 | 0.63 |

†Average correlation to all 5 codon usage measures
nt: nucleotide

**Figure 1.**
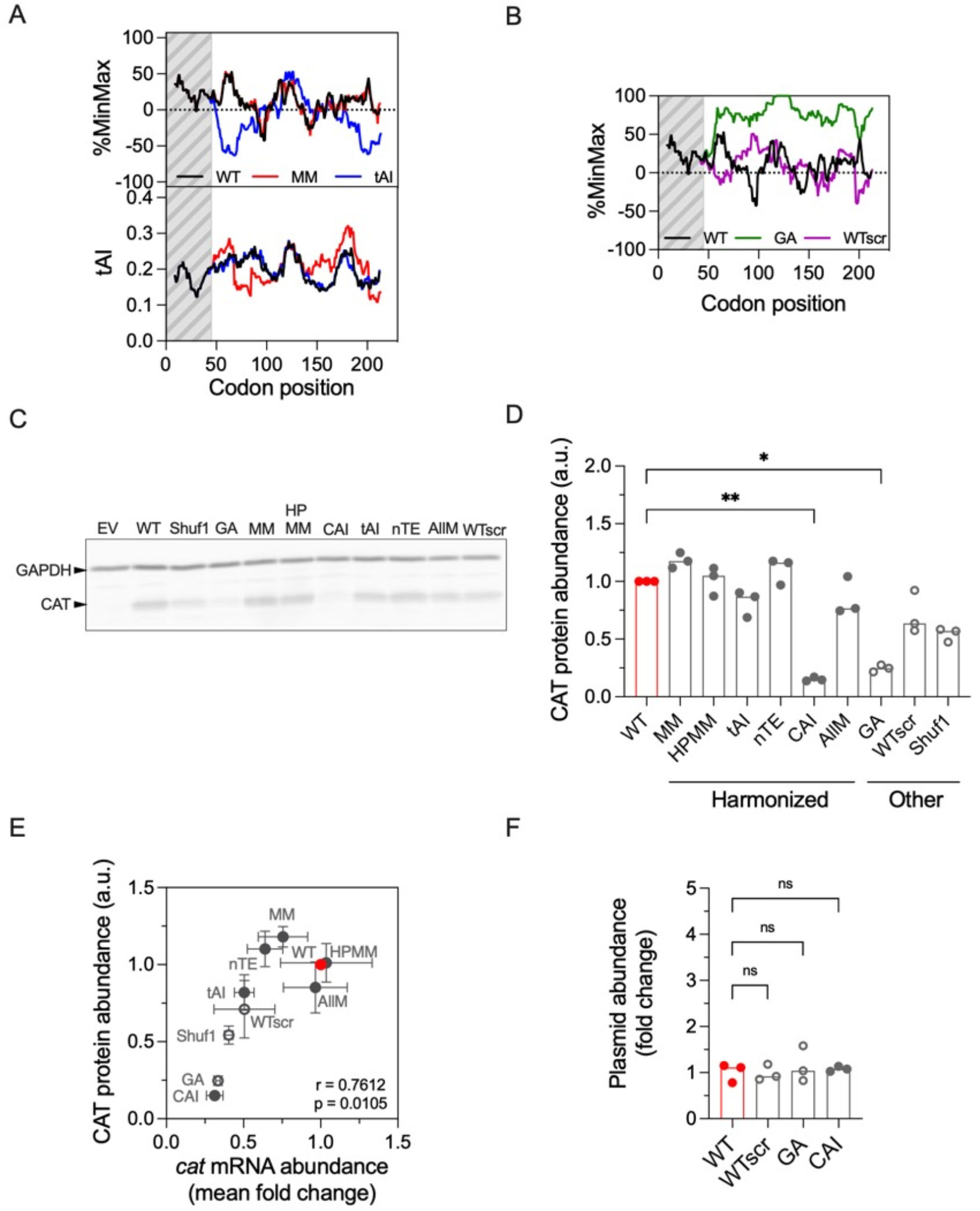
Design and expression of CAT synonymous codon sequences. (**A**) Examples of two harmonized cat mRNA sequences, MM (red) and tAI (blue), generated by matching the codon usage profile of a synonymous mutant sequence to the codon usage profile of WT (black) under a given codon usage measure (top: %MinMax codon usage measure, bottom: tAI codon usage measure) using a sliding window of 17 codons. Harmonized sequences were selected to have low correlation with other codon usage measures. (**B**) In addition to the harmonized cat sequences, two additional synonymous sequences were constructed using other strategies: a sequence optimized by GeneArt (GA) and a scrambled sequence (WTscr) where synonymous codons were swapped but the amino acid sequence was preserved. (**C**) Representative fluorescent western blot to detect synonymous CAT mutant protein abundance in whole cell lysates. EV: empty vector with cat coding sequence removed. GAPDH: loading control. (**D**) Quantification of CAT protein abundance normalized to GAPDH loading control and relative to WT (red bar). Closed circles: harmonized synonymous mutants. Open circles: non-harmonized synonymous mutants. (**E**) Correlation of CAT protein accumulation measured in (D) to the mean fold change of cat mRNA levels relative to WT, determined by RT-qPCR. Significance determined by two-tailed Pearson r correlation with 95% confidence interval. (**F**) Relative plasmid abundance of selected CAT synonymous mutants measured by qPCR. For all categorical scatter plots, bar height indicates the median value of at least three biological replicates (* p < 0.05; ** p < 0.01; Kruskal-Wallis, uncorrected Dunn’s test). Mutants without a label were not significantly different from WT (p > 0.05; Kruskal-Wallis, uncorrected Dunn’s test).

### Synonymous *cat* coding sequences lead to different levels of CAT protein accumulation

To determine whether synonymous recoded *cat* sequences alter CAT protein accumulation, we expressed the ten synonymous *cat* sequences (WT, Shuf1 and the eight new sequences described above) in *E. coli* and quantified intracellular accumulation of CAT protein using fluorescent western blotting. Many of the recoded sequences led to CAT protein levels similar to WT, independent of the codon substitution strategy used (i.e. harmonized vs. other). However, the CAI and GA constructs both led to significantly lower CAT accumulation (**Figure 1C, D**; * p < 0.05; ** p < 0.01, Kruskal-Wallis, uncorrected Dunn’s test). Expression of both sequences also led to dramatically slower *E. coli* growth rates in the presence of chloramphenicol (**Supplementary Figure S1**). We previously showed that when tagged with the C-terminal ssrA degradation sequence, Shuf1-CAT is more susceptible than WT-CAT to degradation by the *E. coli* protease ClpXP (8). To test whether lower CAT protein levels were due to synonymous codon-induced altered protein folding, leading to degradation, we measured *in vivo* protein turnover (**Supplementary Figure S2**) and protein solubility and aggregation (**Supplementary Figure S3**) for all *cat* synonymous mutants. We did not observe significant differences relative to WT, suggesting protein misfolding is not the primary source of the severe defect in protein accumulation for GA and CAI. Similarly, CAT specific activity in cell lysates was also indistinguishable (**Supplementary Figure S4**).

We next tested the effect of synonymous *cat* recoding on *cat* mRNA levels. We measured relative mRNA abundance using RT-qPCR and found a significant correlation of CAT protein abundance to *cat* mRNA levels (**Figure 1E**). To determine whether these different mRNA levels are due to variations in the plasmid copy number between our *cat* constructs, we quantified relative plasmid abundance but found no significant differences in abundance between selected constructs (**Figure 1F**). These data indicate that differences in CAT protein accumulation are primarily due to different levels of *cat* mRNA.

### Synonymous substitutions in cat increase protein accumulation from an upstream coding sequence

In our plasmid, *cat* expression is under the control of the *tet* promoter system, regulated by a negative feedback loop through the *tet* repressor, TetR (**Figure 2A**). We quantified TetR protein levels to determine if differences in *cat* mRNA levels were due to differential expression of *tetR*. We found that TetR protein levels were significantly higher for many recoded *cat* sequences (**Figure 2B**,**C**), leading to an overall inverse relationship between TetR protein accumulation and *cat* mRNA abundance (**Figure 2D**). While the observed decrease in *cat* mRNA levels is consistent with the established model for *tetR*/*tetA* regulation (**Figure 2A**), where higher levels of TetR protein are expected to suppress transcription of *cat* mRNA (33), it was surprising that TetR was unable to repress its own transcription even though it was able to repress *cat*.

**Figure 2.**
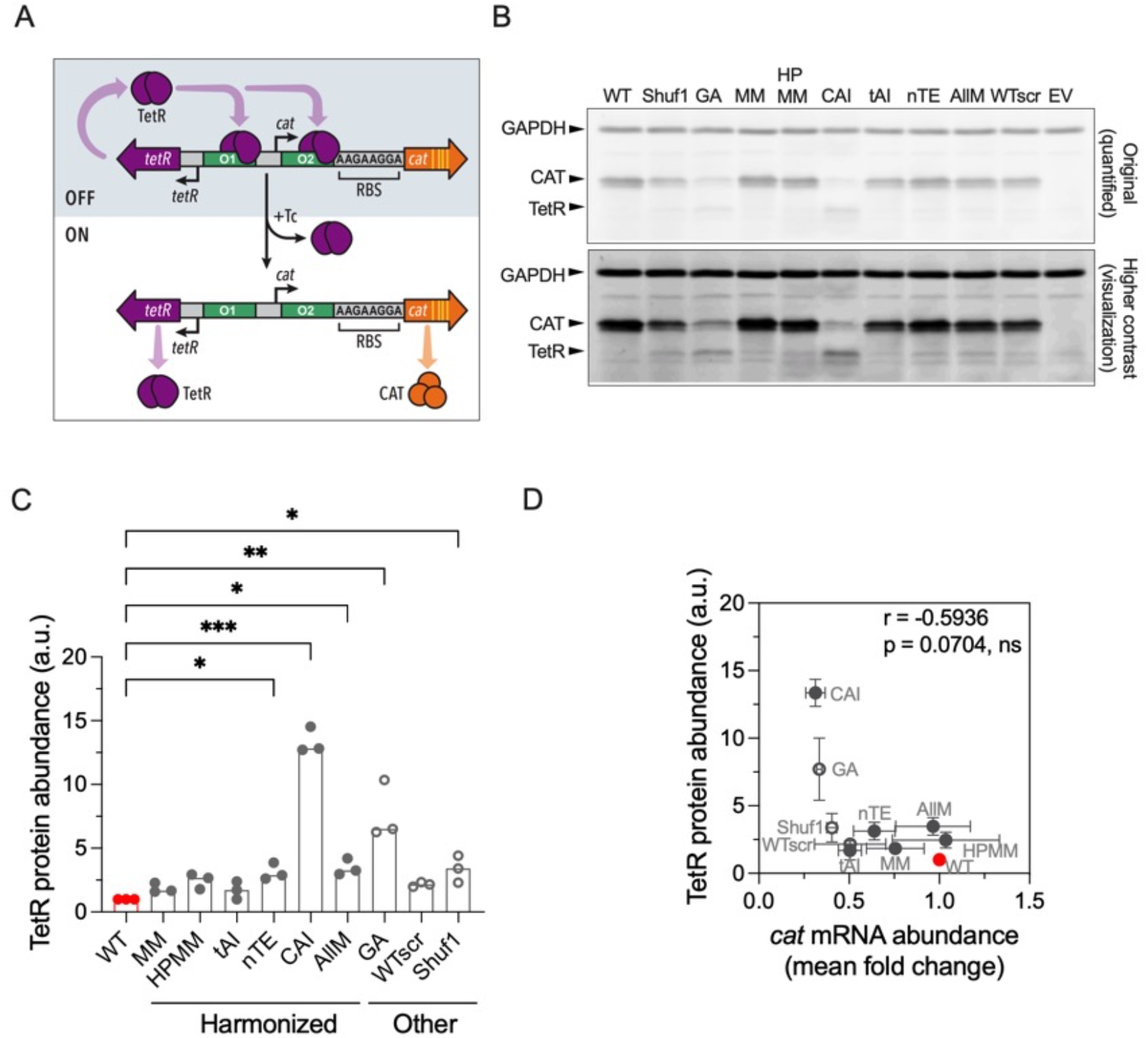
Synonymous codon changes in cat gene increases TetR protein accumulation, which represses cat expression (* p < 0.05; ** p < 0.01; Kruskal-Wallis, uncorrected Dunn’s test). Mutants without a label were not significantly different from WT (p > 0.05; Kruskal-Wallis, uncorrected Dunn’s test). **(A**) Schematic of the overlapping, divergent tet promoter system used to express cat. Both tetR and cat transcription are simultaneously repressed by binding of TetR protein (purple) to the two operator sites (green). Yellow vertical lines indicate location of synonymous codon substitutions in the cat coding sequence, far from the promoter region. (**B**) Representative fluorescent western blot using anti-TetR primary antibody to detect TetR protein in whole cell lysates. Top: blot image used for quantification in (C). Bottom: blot image shown with increased contrast, for display purposes only. EV: empty vector with cat coding sequence removed but tetR gene still present. (**C**) Quantification of TetR protein abundance normalized to loading control, GAPDH, and relative to WT (red bar). Closed circles: harmonized synonymous mutants. Open circles: non-harmonized synonymous mutants. * p < 0.05; ** p < 0.01; *** p < 0.001 (Kruskal-Wallis, uncorrected Dunn’s test). (**D**) TetR protein accumulation has an overall inverse relationship to cat mRNA levels.

To determine the mechanism that causes TetR protein production to be sensitive to *cat* synonymous recoding, we first considered a potential influence from a previously proposed coupling between transcription and translation in bacteria (34). To test if translation elongation of *cat* mRNA could somehow influence *tetR* regulation, we mutated the *cat* ribosome binding site (RBS) to prevent its translation. The elevated accumulation of TetR for GA and CAI persisted (**Figure 3A**), indicating that the connection between *cat* synonymous codon usage and TetR accumulation is independent of *cat* translation.

**Figure 3.**
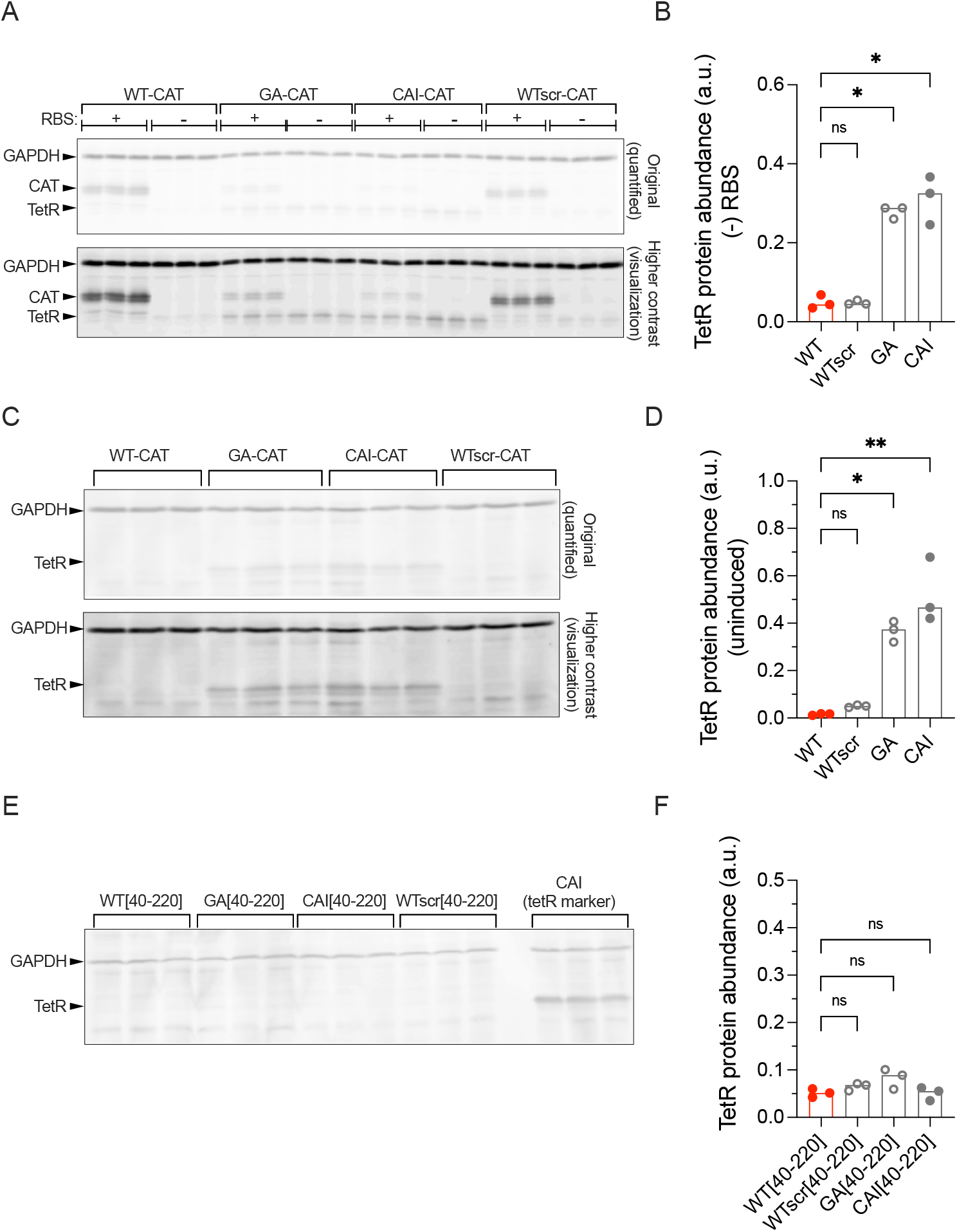
TetR protein upregulation is independent of cat translation and induction but is sensitive to plasmid location of the recoded cat sequence. (**A**) Fluorescent western blot of TetR protein when the cat RBS site is non-functional. (**B**) Quantification of TetR protein when the cat RBS site is non-functional ((-) RBS). * p < 0.05 (Kruskal-Wallis, uncorrected Dunn’s test). (**C**) Fluorescent western blot of TetR protein in the absence of inducer. (**D**) Quantification of TetR protein in the absence of inducer. * p < 0.05; ** p < 0.01 (Kruskal-Wallis, uncorrected Dunn’s test). (**E**) Fluorescent western blot of TetR protein when the synonymous coding region of the selected constructs is cloned into the plasmid backbone of WT-CAT. (**F**) Quantification of TetR protein when the synonymous coding region of the selected constructs is cloned into the plasmid backbone of WT-CAT. TetR abundance was not significantly different from WT; p > 0.05 (Kruskal-Wallis, uncorrected Dunn’s test).

We next considered whether the underlying mechanism leading to increased TetR accumulation is dependent on induction. We found that increased TetR protein accumulation persisted for GA and CAI even in the absence of inducer (**Figure 3B**). A possible mechanism that could lead to increased basal levels of TetR is if the GA and CAI *cat* sequences introduce one or more ectopic TetR binding sites. Conceivably, introduction of these sites could sequester TetR, preventing it from repressing *tetR* transcription but nevertheless blocking efficient transcription of *cat*. To test this mechanism, we inserted the recoded *cat* region (codons 40-220) 287 nucleotides downstream of the *cat* stop codon. We reasoned that if the GA and CAI *cat* sequences harbor additional TetR binding sites, leading to TetR sequestration, this insertion would lead to increased TetR protein accumulation even in the WT *cat* background. However, TetR protein accumulation was similar to WT across all mutant insertions (**Figure 3C**), indicating that the GA and CAI *cat* constructs do not introduce additional TetR binding sites. Taken together, these results indicate that the underlying mechanism leading to increased TetR protein production is independent of regulatory action at the *tet* promoter site.

### TetR is upregulated via cryptic transcription originating within the *cat* coding sequence

An alternative mechanism that could explain increased TetR protein accumulation for recoded *cat* sequences is cryptic transcription originating from within the anti-sense strand of *cat*. Cryptic transcription could result from synonymous codon substitutions if those substitutions inadvertently introduce a new internal promoter and/or modify an enhancer sequence of an existing internal promoter (35, 36). Conceivably, cryptic transcription could bypass the native TetR negative feedback loop and increase the total amount of *tetR* mRNA, leading to increased TetR protein production. To experimentally determine if cryptic transcription is occurring, we designed RT-qPCR primers to amplify *tetR* mRNA that includes the intergenic region upstream of both canonical *tetR* transcription start sites (**Figure 4A, schematic**). We reasoned that amplification of this region should only occur if *tetR* transcription is initiated from an internal promoter within the *cat* coding sequence, on the anti-sense strand. Altogether, four of the nine recoded *cat* sequences (CAI, GA, AllM and Shuf1) led to significant increases in cryptic *tetR* mRNA levels relative to WT *cat*, with the largest increases for the CAI and GA *cat* coding sequences (91-fold and 29-fold more cryptic *tetR* mRNA than WT, respectively) (**Figure 4A**). Surprisingly, TetR protein abundance correlated closely with cryptic *tetR* transcript abundance (**Figure 4B**) but not with total *tetR* mRNA levels (**Figure 4D**). Additionally, there was no correlation between cryptic and total *tetR* mRNA levels (**Figure 5A**), strongly suggesting that the observed increase in TetR protein accumulation is mainly driven by the presence of cryptic *tetR* transcripts.

**Figure 4.**
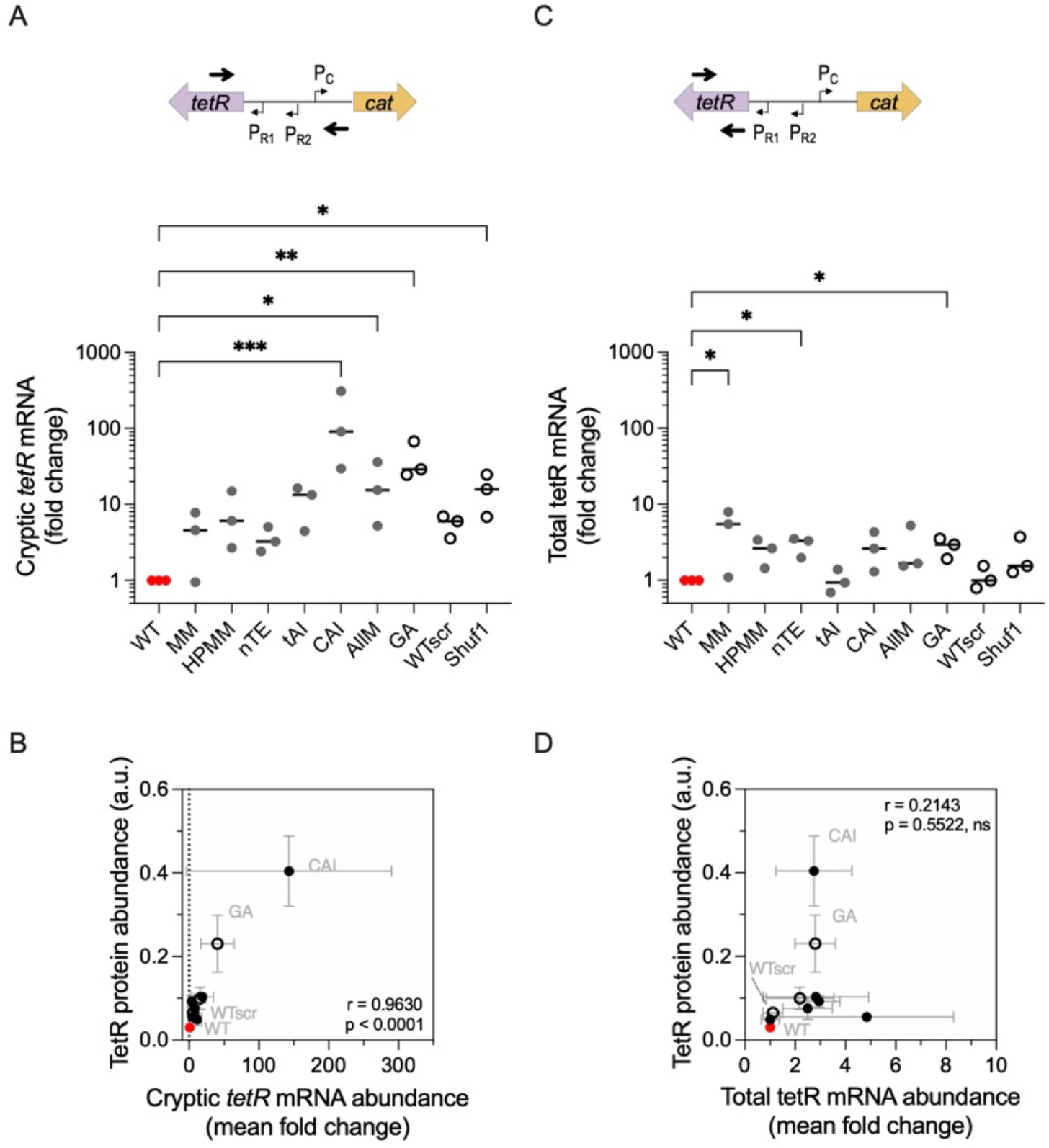
Some cat synonymous substitutions enhance cryptic transcription of tetR (* p < 0.05; ** p < 0.01; *** p < 0.001; Kruskal-Wallis, uncorrected Dunn’s test). (**A**) Analysis of cryptic tetR mRNA abundance relative to WT cat, measured using RT-qPCR. P_R1_ and P_R2_: canonical tetR transcription start sites 1 and 2, respectively. P_C_: cat transcription start site. Bold arrows indicate locations of forward and reverse primer; note reverse primer is designed to bind 41 bp upstream of P_R2_. Lengths are not drawn to scale. * p < 0.05; ** p < 0.01; *** p < 0.001 (Kruskal-Wallis, uncorrected Dunn’s test). (**B**) TetR protein abundance is significantly correlated to cryptic tetR mRNA abundance by two-tailed Pearson r correlation with 95% confidence interval. (**C**) Analysis of total tetR mRNA abundance relative to WT using RT-qPCR. As in panel A, bold arrows indicate locations of forward and reverse primers. * p < 0.05 (Kruskal-Wallis, uncorrected Dunn’s test). Mutants without a label were not significantly different from WT (p > 0.05; Kruskal-Wallis, uncorrected Dunn’s test). (**D**) TetR protein abundance was not significantly correlated to total tetR mRNA abundance by two-tailed Pearson r correlation with 95% confidence interval.

**Figure 5.**
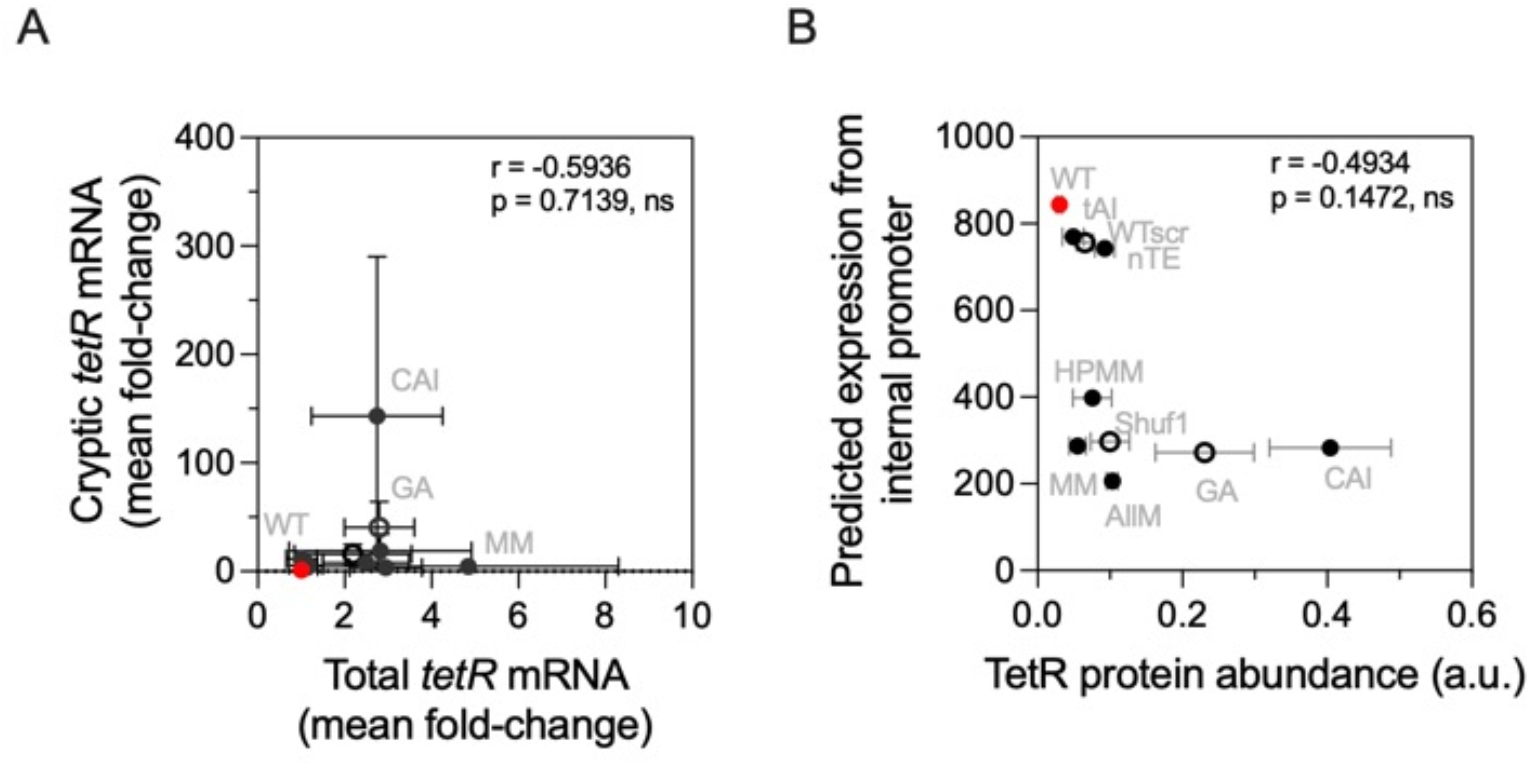
Cryptic tetR transcript levels are not correlated to various features. **(A**) Correlation of cryptic to total tetR transcript levels. There is no significant correlation as determined by two-tailed Pearson r correlation with 95% confidence interval. (**B**) Predicted expression was calculated as described in (38) and was not correlated to TetR protein abundance as determined by two-tailed Pearson r correlation with 95% confidence interval. A higher expression score indicates higher expression levels from predicted promoter.

To determine if the recoded *cat* sequences included a new internal promoter, we analyzed the anti-sense strand of our *cat* synonymous sequences using the bacterial promoter prediction program BPROM, which uses a weighted matrix to detect and score potential promoter sites against the *E. coli* -10 and -35 consensus promoter sequence (37). The similarity of BPROM predicted promoters to the RNAP consensus sequence has previously been shown to be predictive of transcriptional regulation in *E. coli* (38). In total, two potential promoters were identified in the anti-sense strand of all *cat* sequences tested here, including WT. The first predicted promoter resides in the unaltered 5’ region of the *cat* coding sequence; thus, it is identical in all constructs and is not considered further here. Each sequence also contains a second predicted promoter toward the 3’ end of the *cat* gene (**Table 2**). For both WT and WTscr, this 3’ predicted promoter is in the same location and closely matches the consensus RNAP binding sequence. In contrast, although both CAI and GA mutant sequences also have a predicted promoter in a similar location, it is less similar to the RNAP consensus binding sequence. Hence, based on RNAP consensus similarity alone, the WT and WTscr *cat* sequences are predicted to result in more *tetR* transcription than GA and CAI, not less. Likewise, using the BPROM output to calculate expected TetR expression from each predicted internal promoter (as previously described (38)), led to a predicted trend in TetR protein levels that is opposite from the measured TetR abundance levels (**Figure 5B**). Taken together, the inconsistencies between the computational predictions and measured tetR mRNA and TetR protein levels suggests that (i) there may be additional factors that regulate transcription from the predicted internal promoter site within *cat*, or (ii) the internal promoter identified by BPROM does not produce the cryptic *tetR* mRNA transcripts we detected by RT-qPCR.

**Table 2.**
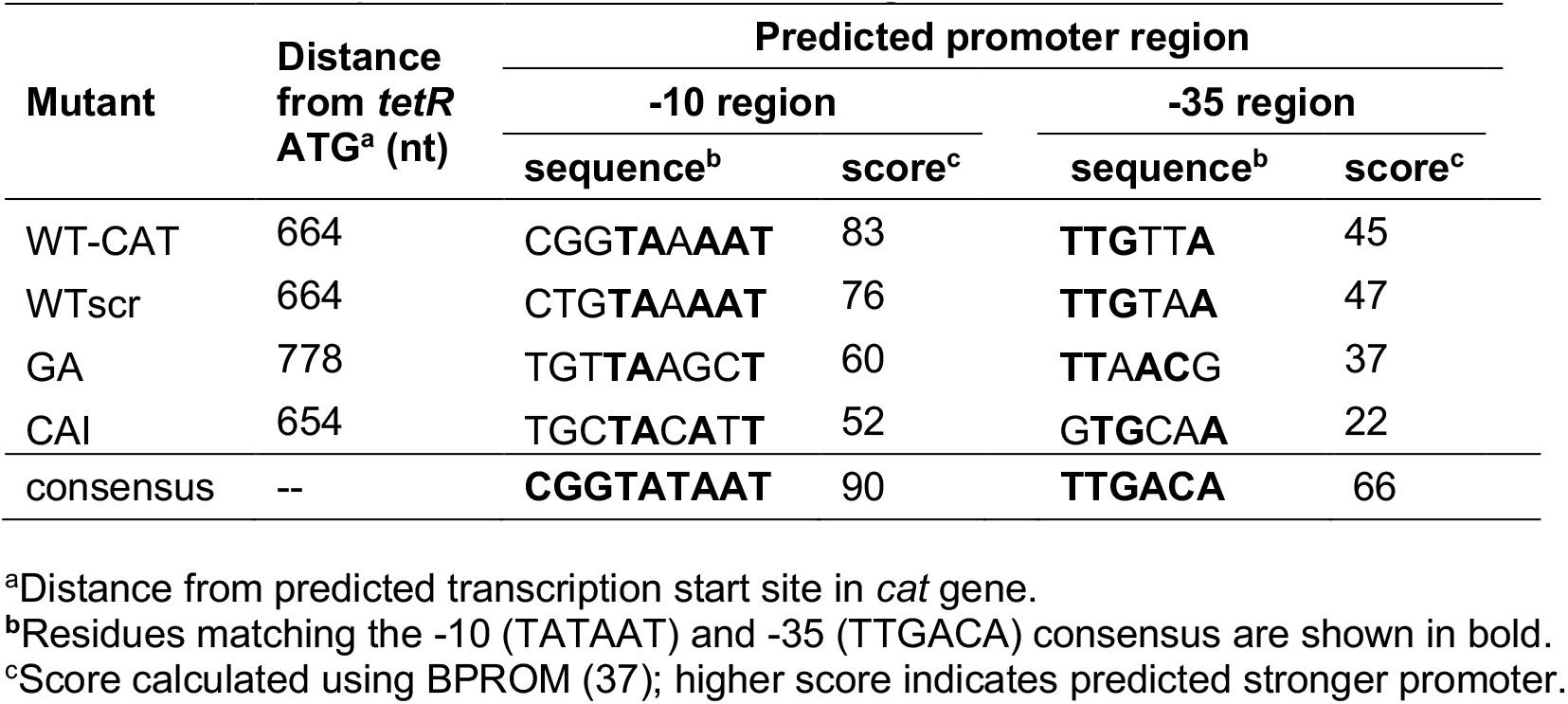
Predicted promoter location and strength.

## DISCUSSION

A growing body of evidence has demonstrated that synonymous codon substitutions can modulate a wide range of mechanisms that affect expression of the encoded gene, including its transcriptional level (3, 20, 39), translational efficiency (4, 29, 31) and co-translational folding (7, 8, 40). Here we show that in addition to these well-characterized intragenic effects, synonymous codon substitutions can also play a major role in regulating transcription and translation of a gene located upstream of the recoded coding sequence, via enhanced cryptic transcription. In our system, this cryptic transcription led to marked phenotypic consequences because the upstream gene is the transcription factor TetR, which regulates expression of a selectable marker (resistance to chloramphenicol) (**Figure 6**). One could imagine that if the upstream gene instead encoded a more general transcription factor, cryptic transcription could lead to widespread, coordinated regulation of multiple genes located elsewhere on a plasmid or genome. In this way, cryptic transcription may provide an explanation for currently poorly understood global fitness effects observed upon synonymous codon substitutions, especially for substitutions located in genomic regions rich in directional and/or divergent promoters (41). Of note, the effects of *cat* codon substitutions on *tetR* transcription and translation were significantly larger than any effect on CAT protein folding efficiency, demonstrating that transcriptional regulation can be more sensitive than co-translational folding to synonymous codon usage, even in a system with demonstrated effects of synonymous codons on folding efficiency (8). Similarly, others have shown that intragenic 5’ codon substitutions typically result in larger effects on translational efficiency of the encoded protein than its co-translational folding (4, 29, 42).

**Figure 6.**
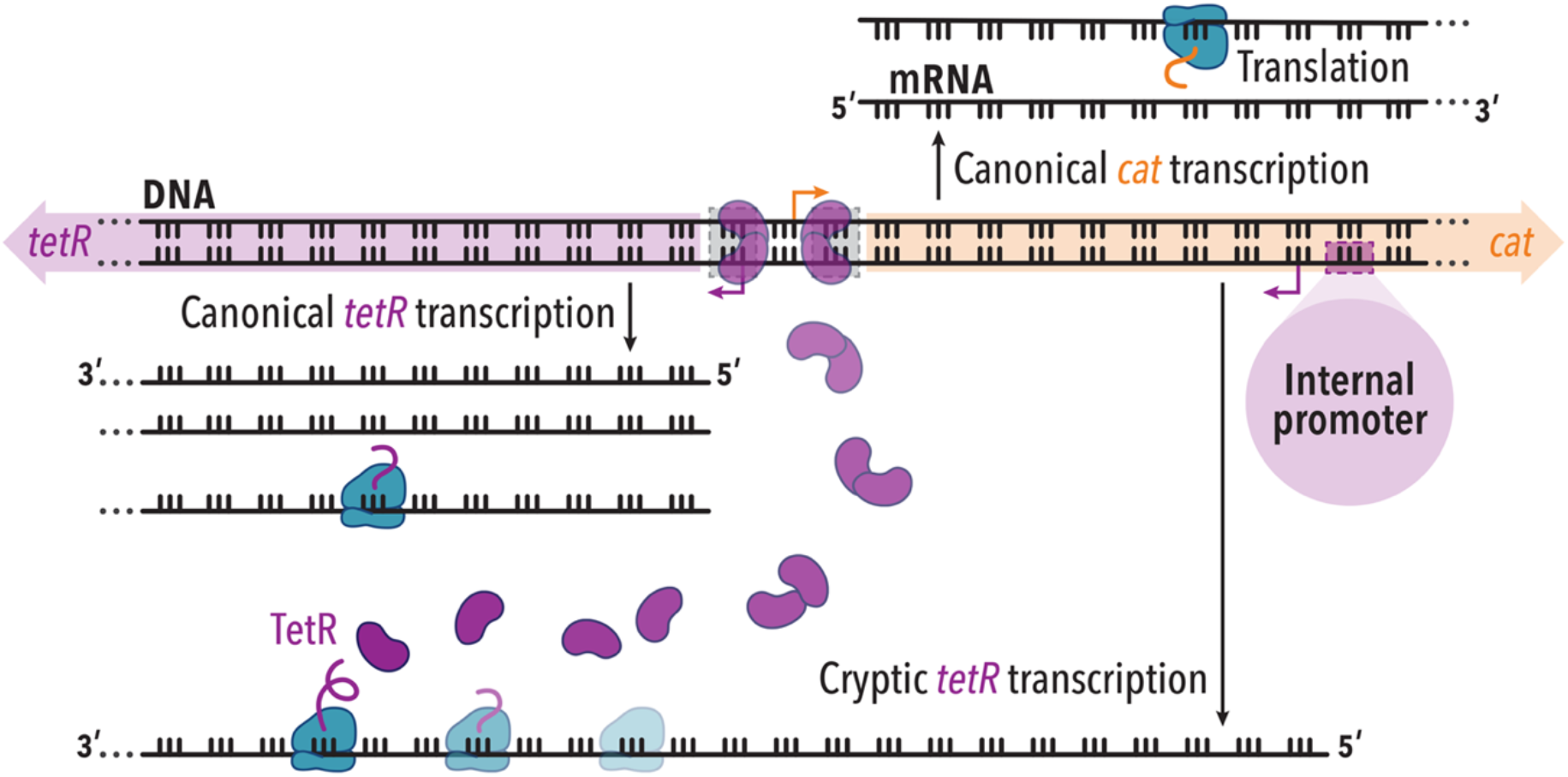
Model for synonymous substitutions upregulating transcription and translation of an upstream gene. Synonymous substitutions in the cat coding sequence (orange) enhance transcription from an internal promoter on the antisense strand. Although this cryptic transcription is a subset of total tetR mRNA, it correlates closely with the resulting level of TetR protein. The correlation between cryptic RNA abundance and TetR protein level may occur because the cryptic RNA is translated with higher efficiency than mRNA from the canonical tetR transcription start site, as shown here. Alternatively (not shown), the cryptic RNA may act as a non-coding RNA to upregulate translation of the canonical tetR RNA. Regardless of the precise mechanism, cryptic transcription of tetR provides a mechanism to maintain TetR-mediated transcriptional repression of cat while bypassing repression of tetR.

There are >10^80^ synonymous sequences that can encode the wild type CAT protein sequence. Of the nine synonymous *cat* sequences studied here, four showed significantly increased levels of cryptic transcription relative to WT. The frequency of these observed changes in transcription demonstrates that sequences that lead to cryptic transcription are readily accessible in synonymous sequence space. These results have two important implications: First, synonymous codon usage may be subject to selective pressures to avoid introducing or enhancing cryptic transcription start sites in protein-coding sequences. In addition to the triggering of regulatory mechanisms as described here, other reasons to avoid internal promoters include avoiding transcriptional interference and the metabolic costs and resulting fitness costs of uncontrolled transcription (38, 43). Second, these selection pressures may be gene context dependent, as in some contexts, selection for sequences that upregulate neighboring genes might provide an advantage (41, 44, 45). More investigation is needed to determine which, if any, genes have evolved to exploit internal promoter sites to regulate neighboring genes.

Of note, while promoter prediction algorithms accurately predict expression level for naturally occurring promoters (38), we did not observe the predicted correlation between predicted expression level and TetR protein abundance. This lack of correlation may complicate computational identification of the impact of synonymous mutations on cryptic transcription levels using existing algorithms. Nevertheless, if selection for or against internal promoter-guided transcription is widespread, it may partially explain why synonymous codon usage is under selection in bacterial genomes (46, 47)(48). A major implication of these findings is that the transcriptional regulatory network and the pressures it places on synonymous codon usage may be significantly larger than our current understanding, which is largely confined to promoters that reside in intergenic regions, outside of coding sequences. Such selective pressures could provide an explanation for pervasive transcription, a term used to describe the widespread detection in bacteria of RNAs derived from unexpected start sites, often on the non-coding strand (43, 49). Our results support the model that these RNAs may arise not merely from random transcriptional noise but instead could have important functional roles.

Perhaps most surprisingly, TetR protein accumulation was well correlated with the subset of *tetR* mRNA arising from the internal promoter site rather than total *tetR* mRNA, implying that the cryptic *tetR* mRNA leads to more efficient TetR translation than *tetR* mRNA produced from the canonical transcription start site. More investigation is needed to determine the mechanistic origin of cryptic *tetR* mRNA on TetR protein accumulation. For example, the longer 5’ UTR could modulate *tetR* mRNA structure, improving access to the ribosome binding site. Alternatively, the cryptic RNA may not itself be translated, but instead regulate as a noncoding RNA translation of *tetR* mRNA transcribed from the canonical transcription start sites. Regardless of the precise mechanism, if cryptic transcription of neighboring genes leads to mRNA with higher translational efficiency, cryptic transcription may contribute to the overall low correlation between total mRNA and protein abundance observed in *E. coli* and other organisms (50–52).

## SUPPLEMENTARY DATA

Supplementary Figures, Tables and Methods are available.

## ACKNOWLEDGEMENTS

The authors are grateful to Allen Buskirk, Patricia Champion and Qing Luan for helpful discussions and Jacob Diehl for assistance with some experiments.

