## Supplementary Methods, Figures & Tables for "Synonymous codon substitutions regulate transcription and translation of an upstream gene"

*Synonymous cat coding sequences*

*WT-CAT*

*ATGCATCACCATCACCATCACCATAACTATACAAAATTTGATGTAAAAAATTGGGTTCGCCGTGAGCA
TTTTGAGTTTTATCGGCATCGTTTACCATGTGGTTTTAGCTTAACAAGCAAAATTGATATCACGACGTT
AAAAAAGTCATTGGATGATTCAGCGTATAAGTTTTATCCGGTAATGATCTATCTGATTGCTCAGGCCG
TGAATCAATTTGATGAGTTGAGAATGGCGATAAAAGATGATGAATTGATCGTATGGGATTCAGTCGAC
CCACAATTCACCGTATTCCATCAAGAAACAGAGACATTTTCAGCACTGAGTTGCCCATACTCATCCGA
TATTGATCAATTTATGGTGAATTATTTATCGGTAATGGAACGTTATAAAAGTGATACCAAGTTATTTCC
TCAAGGGGTAACACCAGAAAATCATTTAAATATTTCAGCATTACCTTGGGTTAATTTTGATAGCTTTAA
TTTAAATGTTGCTAATTTTACCGATTATTTTGCACCCATTATAACAATGGCAAAATATCAGCAAGAAGG
GGATAGACTGTTATTGCCGCTCTCAGTACAGGTTCATCATGCAGTTTGTGATGGCTTCCATGTTGCAC
GCTTTATTAATCGGCTACAAGAGTTGTGTAACAGTAAATTAAAATAA*

*%MinMax (MM-CAT)*

*ATGCATCACCATCACCATCACCATAACTATACAAAATTTGATGTAAAAAATTGGGTTCGCCGTGAGCA
TTTTGAGTTTTATCGGCATCGTTTACCATGTGGTTTTAGCTTAACAAGCAAAATTGATATCACGACGTT
AAAAAAGTCATTGGATGATAGCGCTTACAAATTCTACCCGGTCATGATCTATCTGATTGCTCAGGCCG
TGAATCAATTTGACGAACTACGAATGGCGATAAAAGATGATGAATTAATCGTCTGGGACAGTGTTGAT
CCTCAATTCACTGTGTTCCACCAAGAAACTGAGACTTTCAGCGCGTTAAGCTGCCCGTATTCCTCCG
ATATCGATCAATTTATGGTGAACTACTTAAGCGTGATGGAGCGTTATAAAAGCGATACTAAACTATTC
CCCCAAGGAGTTACGCCCGAAAATCATCTAAACATCTCGGCCCTCCCGTGGGTAAACTTTGACAGCT
TTAACTTAAACGTCGCTAACTTTACCGATTATTTCGCTCCCATCATCACAATGGCAAAATACCAACAG
GAAGGGGACCGTTTGCTGCTACCGTTGAGCGTACAGGTACATCACGCGGTCTGCGACGGCTTTCAT
GTGGCCCGTTTCATCAATCGACTCCAAGAGTTATGTAATTCTAAGCTGAAATAA*

*High-Phi %MinMax (HPMM-CAT)*

*ATGCATCACCATCACCATCACCATAACTATACAAAATTTGATGTAAAAAATTGGGTTCGCCGTGAGCA
TTTTGAGTTTTATCGGCATCGTTTACCATGTGGTTTTAGCTTAACAAGCAAAATTGATATCACGACGTT
AAAGAAGTCATTGGACGATTCTGCATATAAATTTTATCCTGTAATGATCTACCTTATAGCGCAGGCAG
TAAATCAATTCGACGAGTTACGAATGGCCATTAAGGACGACGAGTTGATAGTTTGGGACTCCGTTGA
CCCACAATTCACGGTTTTCCATCAAGAGACCGAGACCTTTTCTGCTCTCTCATGCCCCTATAGCTCAG
ACATAGACCAATTTATGGTCAATTACCTCTCGGTCATGGAGCGCTACAAAAGCGATACTAAGCTTTTT
CCACAAGGTGTCACGCCAGAGAATCATTTAAATATAAGCGCATTACCCTGGGTGAATTTCGATTCGTT
TAATCTAAATGTAGCCAATTTCACAGATTATTTTGCCCCAATCATCACCATGGCCAAGTATCAACAAGA
AGGAGACCGTTTATTACTTCCGCTGAGTGTGCAGGTTCATCATGCAGTATGTGACGGATTTCACGTA
GCGCGATTCATTAATCGATTACAGGAGCTCTGTAATTCCAAGCTGAAGTAA*

*Codon Adaptation Index (CAI-CAT)*

*ATGCATCACCATCACCATCACCATAACTATACAAAATTTGATGTAAAAAATTGGGTTCGCCGTGAGCA
TTTTGAGTTTTATCGGCATCGTTTACCATGTGGTTTTAGCTTAACAAGCAAAATTGATATCACGACGTT
AAAAAAGTCATTAGACGATTCAGCTTACAAATTTTATCCGGTTATGATTTACTTAATTGCTCAGGCCGT
GAACCAATTCGATGAGCTACGAATGGCCATAAAGGATGACGAGCTGATTGTCTGGGATTCAGTTGAT
CCGCAGTTTACAGTGTTCCACCAGGAGACAGAAACATTTTCTGCTCTTAGTTGTCCGTATAGTAGTGA
TATTGACCAATTCATGGTTAATTACCTCTCCGTAATGGAACGGTACAAGTCGGATACGAAACTTTTCC
CACAGGGGGTAACGCCTGAGAATCACCTTAACATATCGGCCCTCCCCTGGGTAAACTTCGACTCGTT
TAATCTAAATGTAGCAAATTTCACTGATTACTTTGCACCCATCATAACGATGGCCAAGTATCAACAAGA
GGGTGACCGACTCCTTTTACCGCTCTCAGTTCAAGTACATCACGCCGTATGCGACGGTTTCCACGTA
GCTCGGTTCATCAATCGACTACAAGAACTCTGCAACTCGAAATTAAAGTAA*

*tRNA Adaptation Index (tAI-CAT)*

*ATGCATCACCATCACCATCACCATAACTATACAAAATTTGATGTAAAAAATTGGGTTCGCCGTGAGCA TTTTGAGTTTTATCGGCATCGTTTACCATGTGGTTTTAGCTTAACAAGCAAAATTGATATCACGACGTT AAAAAAGTCTCTTGATGACTCTGCCTATAAGTTCTATCCTGTAATGATATACCTCATAGCACAAGCGG TTAACCAATTCGACGAGCTTAGAATGGCCATTAAGGATGACGAACTAATTGTATGGGACTCCGTCGA CCCTCAGTTCACTGTATTCCATCAGGAAACCGAAACATTCAGTGCCCTATCCTGTCCATATTCGTCGG ATATCGACCAGTTTATGGTTAACTATCTGTCAGTTATGGAAAGATATAAAAGTGACACCAAACTTTTCC CGCAGGGTGTGACTCCTGAGAATCACCTCAACATTTCCGCTTTACCATGGGTGAATTTTGATTCTTTT AATCTTAATGTGGCTAATTTTACAGATTATTTCGCCCCGATCATAACTATGGCAAAGTATCAGCAAGA GGGCGATCGCTTGCTCCTCCCTTTGTCTGTCCAAGTACACCACGCTGTTTGCGACGGATTTCATGTA GCCAGGTTTATAAACAGATTACAGGAGCTTTGTAATTCCAAGTTGAAATAA*

*Normalized Translation Efficiency (nTE-CAT)*

*ATGCATCACCATCACCATCACCATAACTATACAAAATTTGATGTAAAAAATTGGGTTCGCCGTGAGCA TTTTGAGTTTTATCGGCATCGTTTACCATGTGGTTTTAGCTTAACAAGCAAAATTGATATCACGACGTT AAAAAAATCCTTAGACGATAGTGCGTATAAGTTTTATCCAGTGATGATCTACCTTATCGCACAGGCTG TTAACCAATTTGATGAGCTGCGCATGGCCATAAAAGATGACGAGCTGATCGTATGGGATTCGGTAGA CCCTCAATTCACCGTATTCCACCAAGAAACAGAAACGTTCTCGGCACTCTCGTGTCCGTACTCAAGT GATATCGACCAATTTATGGTGAATTATCTTTCTGTAATGGAGCGGTATAAGTCTGATACCAAGCTATT CCCTCAAGGTGTCACACCTGAAAATCACTTGAATATCAGTGCTCTCCCATGGGTTAACTTTGATTCCT TTAACCTGAACGTTGCTAATTTCACCGACTACTTCGCTCCGATTATAACAATGGCCAAGTATCAACAA GAGGGTGACCGGTTGCTTTTACCACTTTCCGTGCAAGTGCACCACGCCGTTTGTGATGGTTTTCATG TGGCTCGGTTCATAAATCGGCTCCAGGAGTTGTGTAACTCTAAACTGAAATAA*

*AllModel (AllM-CAT)*
*ATGCATCACCATCACCATCACCATAACTATACAAAATTTGATGTAAAAAATTGGGTTCGCCGTGAGCA TTTTGAGTTTTATCGGCATCGTTTACCATGTGGTTTTAGCTTAACAAGCAAAATTGATATCACGACGTT AAAAAAGTCACTAGATGATTCGGCGTATAAATTCTATCCAGTGATGATTTATTTAATCGCCCAGGCCG TGAATCAGTTTGACGAGCTCAGGATGGCCATAAAGGATGATGAACTCATCGTGTGGGACAGTGTTGA CCCTCAATTCACGGTCTTCCATCAGGAGACAGAAACTTTTTCCGCATTGTCGTGTCCATACTCCTCAG ATATTGATCAGTTTATGGTCAATTATCTCAGCGTTATGGAGCGCTATAAATCAGACACAAAACTATTTC CACAAGGGGTTACTCCCGAGAATCATCTAAATATTTCCGCACTACCGTGGGTGAATTTTGATAGTTTT AATCTAAATGTCGCCAACTTCACAGATTATTTTGCCCCCATAATCACTATGGCGAAGTACCAACAAGA AGGTGATAGGTTGTTGTTACCACTGAGTGTTCAGGTCCACCACGCTGTGTGTGATGGCTTTCATGTA GCACGGTTTATAAACAGGCTACAGGAATTATGTAATTCTAAGTTAAAGTAA*

*GeneArt Optimization (GA-CAT)*

*ATGCATCACCATCACCATCACCATAACTATACAAAATTTGATGTAAAAAATTGGGTTCGCCGTGAGCA TTTTGAGTTTTATCGGCATCGTTTACCATGTGGTTTTAGCTTAACAAGCAAAATTGATATCACGACGTT AAAAAAGTCATTGGATGATAGCGCCTATAAATTCTATCCGGTGATGATTTATCTGATTGCCCAGGCAG TTAACCAGTTTGATGAACTGCGTATGGCCATCAAAGATGATGAGCTGATTGTTTGGGATAGCGTTGAT CCGCAGTTTACCGTTTTTCATCAAGAAACCGAAACCTTTAGCGCACTGAGCTGTCCGTATAGCAGCG ATATTGATCAGTTTATGGTGAACTATCTGAGCGTGATGGAACGCTATAAAAGCGATACCAAACTGTTT CCGCAGGGTGTTACACCGGAAAATCATCTGAATATTTCAGCACTGCCGTGGGTGAACTTTGATTCCT TTAATCTGAATGTGGCCAACTTCACCGATTATTTTGCTCCGATTATTACCATGGCCAAATATCAGCAA GAAGGTGATCGTCTGCTGCTGCCGCTGAGCGTTCAGGTTCATCATGCAGTTTGTGATGGTTTTCATG TTGCCCGTTTTATCAATCGTCTGCAAGAACTGTGTAACAGCAAACTGAAATAA*

*WTscramble (WTscr-CAT)*

*ATGCATCACCATCACCATCACCATAACTATACAAAATTTGATGTAAAAAATTGGGTTCGCCGTGAGCA TTTTGAGTTTTATCGGCATCGTTTACCATGTGGTTTTAGCTTAACAAGCAAAATTGATATCACGACGTT AAAAAAGTCATTGGACGATAGCGCATACAAATTCTATCCAGTGATGATTTATCTAATAGCACAAGCAG*

*TCAACCAGTTCGATGAATTACGTATGGCAATTAAGGATGATGAGCTCATTGTGTGGGATTCGGTTGAT CCGCAGTTTACAGTTTTTCATCAGGAGACCGAAACCTTCTCCGCGTTATCATGTCCGTATAGTTCAGA TATCGATCAATTTATGGTAAATTATCTGTCAGTTATGGAGCGGTATAAGTCAGATACAAAATTGTTTCC CCAAGGCGTTACCCCTGAAAATCATTTGAATATCAGTGCGTTGCCATGGGTAAATTTTGATTCATTTA ATTTGAATGTAGCAAATTTTACAGATTATTTTGCCCCAATAATTACAATGGCTAAATATCAACAAGAAG GGGATCGCTTACTGCTGCCTTTAAGTGTTCAAGTACATCATGCTGTATGCGATGGGTTTCATGTAGCA AGATTTATTAATAGATTACAAGAATTATGTAATTCAAAATTAAAATAA*

*Shuf1-CAT*

*ATGCATCACCATCACCATCACCATAACTATACAAAATTTGATGTAAAAAATTGGGTTCGCCGTGAGCA TTTTGAGTTTTATCGGCATCGTTTACCATGTGGTTTTAGCTTAACAAGCAAAATTGATATCACGACGTT AAAAAAGTCATTGGATGATTCAGCGTATAAGTTTTACCCCGTGATGATATACTTAATTGCCCAAGCAG TTAACCAATTTGACGAGCTCCGAATGGCGATTAAAGATGATGAACTCATTGTGTGGGACTCCGTCGA TCCGCAATTTACTGTATTCCATCAGGAGACTGAAACTTTTAGTGCGCTATCCTGCCCCTATAGCTCGG ACATCGATCAATTTATGGTGAATTACCTGTCGGTCATGGAAAGGTATAAGTCCGACACAAAATTATTT CCCCAGGGCGTGACGCCAGAAAACCATCTAAACATATCCGCGCTGCCATGGGTTAACTTTGACTCGT TCAACTTAAATGTAGCTAACTTTACTGACTATTTTGCGCCGATCATAACCATGGCCAAATACCAACAG GAAGGGGATAGGCTCTTGTTACCCTTGAGCGTCCAGGTGCATCACGCCGTGTGTGACGGCTTTCAT GTTGCACGCTTCATTAACAGATTACAGGAGTTATGTAATTCTAAATTGAAGTAA*

*Codon scores used in CHARMING algorithm to generate harmonized synonymous sequences*

All codon scores are specific to *E. coli* and are listed in **Table S4**. Previously published codon score values were used for Codon Adaptation Index (CAI) (1), tRNA adaptation index (tAI) (2), codon usage frequencies (3) (%MinMax), and computationally determined codon frequencies at high gene expression levels defined as log_10(phi) = 0.75 (High-Phi) in the ROC-SEMPR model described in (4).

*E. coli* nTE values were calculated following the previously published methodology for *S. cerevisiae* (5). TE is calculated as: tAI/(weighted codon usage). The tAI values used here are the same codon scores described in the previous paragraph. Weighted codon usage was calculated by counting how many times a codon appears in each annotated gene multiplied by mRNA counts (fpkm) of that gene. mRNA counts were retrieved from NCBI GEO database, series GSE58325, sample GSM1406483. TE codon scores were normalized to 1 to generate nTE scores. The gene CAI ($CAI_g$) values were calculated using the web tool from Biologics International Corp: https://www.biologicscorp.com/tools/CAICalculator#.YRaJuVNKhTY.

*Growth assay*

Saturated overnight cultures of *E. coli* KA12 were prepared by 5 mL of Luria Broth (LB) containing 100 ug/mL ampicillin in 14 mL round-bottom Falcon tubes with a single colony from a freshly streaked plate, followed by growth with shaking at 37°C for 16 h. Then, 12-well plates containing 3 mL LB, 100 µg/mL ampicillin, 600 ng/mL tetracycline (tc) to induce gene expression, and 350 µg/mL chloramphenicol (cam) to challenge bacterial growth, were inoculated to $OD_{600}$ =0.05. Cells were grown with continuous double-orbital shaking at 37°C for 12 h in a Synergy H1 microplate reader (BioTek), recording $OD_{600}$ every 10 min. Time to maximum growth rate is the time corresponding to the maximum of the first derivative of the growth curve.

*Protein solubility using fluorescent western blotting*

Three biological replicates of *E. coli* KA12 were grown as described above. After 3 hours of growth and induction, a 2 mL aliquot of the culture was pelleted at 4,600x*g* for 10 min at 4°C. The supernatant was discarded and cell pellets were frozen at -80°C. Cell pellets were thawed and resuspended to $OD_{600}$ =10 using lysis buffer (100 mM Tris pH 7.5, 1 mM PMSF, 5 mM EDTA). Cells were lysed by six freeze/thaw cycles using liquid nitrogen for freezing and a room temperature water bath for thawing. After the last thaw cycle, cells were treated with 20 U/mL DNase (RNase free) (Invitrogen) and supplemented with 25 mM $MgSO_4$ at 4°C for 30 min. Treated lysates were centrifuged at 15,000 rpm for

15 min at 4°C and the cleared cell lysate was transferred to a clean 1.5 mL microcentrifuge tube. Cleared cell lysates and pellets were stored at -80°C. Stored samples were thawed at room temperature. Pellets were prepared for fluorescent western blotting as described above. Cleared lysates were prepared by mixing 20 μL lysate, 10 uL SDS loading dye, 5 mM βME, and 5 mM EDTA.

*Protein turnover in vivo with retapamulin*

To determine CAT susceptibility to degradation *in vivo*, conditions of similar CAT protein production were determined for all mutants by adjusting the tetracycline inducer concentration as follows:

| mutant | tc (ng/mL) |
|---|---|
| WT | 200 |
| Shuf1 | 600 |
| GA | 1200 |
| MM | 250 |
| HPMM | 300 |
| HPCAI | 1300 |
| tAI | 400 |
| nTE | 300 |
| AllM | 400 |
| scr | 400 |

KA12 transformed with plasmids containing CAT were grown in 12-well plates containing 3 mL of LB with 100 μg/mL ampicillin (amp) and variable amounts of tetracycline (tc) for 3 hours at 37°C with double orbital shaking. To remove tc from the media, cells were spun at 4,600 x g, room temperature, for 10 min and the supernatant was discarded. Pellets were rinsed with LB and spun down again. Pellets were resuspended in 3 mL of fresh LB containing 100 μg/mL ampicillin and transferred to a new 12-well plate. A 300 μL aliquot was transferred to a 1.5 mL tube (t=0) and spun down at 4,600 xg, 4°C, for 10 min. The remaining cells in the 12-well plate were treated with 200 μg/mL of retapamulin (Ret) to inhibit translation initiation. Every hour for four hours, a 300 μL aliquot was spun down as described above and all pellets were frozen at -80°C. Frozen cell pellets were then thawed at room temperature and prepared for SDS-PAGE followed by fluorescent western blotting as described above.

*CAT specific activity measurements*

Acetylation of chloramphenicol was quantified in the following manner. All reagents and solvents were LC-MS grade and obtained from Sigma (St. Louis, MO) unless otherwise stated. Three biological replicates of *E. coli* KA12 were grown as described above. After 3 hours of growth and induction, a 2 mL aliquot of the culture was pelleted at 4,600 xg for 10 min at 4°C. The supernatant was discarded and cell pellets were frozen at -80°C. Cell pellets were thawed and resuspended to $OD_{600}$ 10.0 in lysis buffer (100 mM Tris pH 7.5, 5 mM EDTA). Cells were lysed by six freeze/thaw cycles using liquid nitrogen for freezing and a room temperature water bath for thawing. After the last thaw cycle, lysates were centrifuged at 15,000 rpm for 15 min at 4°C and 120 μL of the cleared cell lysate was transferred to a clean 1.5 mL microcentrifuge tube. Enzymatic activity of CAT was performed by addition of 5 μL of 5 mg/mL acetyl-CoA and 2 μL of 3.4 mg/mL CAM. The reaction mixture was incubated at 37ºC for 3 hours and quenched by addition of formic acid (FA) to a final concentration of 1.2%. 60 mM (20.8 μL) d5-CAM (Santa Cruz Biotechnology, Santa Cruz, CA) was added as a stable-isotope internal standard. CAM was extracted from the reaction mixture by 2:1 addition of ethyl acetate. The organic fraction was collected and dried down by speed-vac (ThermoFisher) and stored at -20°C until analysis. The extracted CAM was reconstituted in 100 μL 15% acetonitrile (ACN), 0.1% FA and vortexed extensively. Reconstituted CAM was further diluted ten-fold in the same solvent and 10 μL was analyzed in triplicate via LC-MS. Reverse-phase μHPLC-MS was performed using a Ultimate2000 (Dionex) with a HALO 160Å $C_{18}$ column (Advanced Materials Technology) with a 10 min linear gradient from 15-100% ACN in 0.1% FA. Mass spectra were collected on a Bruker Impact II in the negative ion mode with a mass range of 150-3000 m/z, at 1 spectrum/s, an ESI voltage of -1800 V, and gas flow of 7 L/min at 200ºC. The peak areas for the extracted ion chromatogram of CAM ($[M-H]^-$, m/z 321.0076 ± 0.01) and d5-CAM ($[M-H]^-$, m/z 326.0387 ± 0.01) were integrated using DataAnalysis (BrukerDaltonik). The CAM peak area was normalized to the d5-CAM internal standard and amount of acetylated chloramphenicol was determined by the difference between samples from cells expressing empty vector and cells expressing a *cat* coding sequence.

In parallel, CAT protein abundance was quantified in the following manner. First, 14 µg of the clarified cell lysate prepared above was reduced with TCEP at 95ºC for 5 min and alkylated with iodoacetic acid for 30 min as previously described (6). The lysate was loaded onto a S-Trap micro (Profiti) and washed following the manufacturer's protocol, then digested in 67 µL (100 mM) TEAB and 1 µg trypsin overnight at 37ºC. The peptides were dried down and desalted using ZipTips (Millipore) following the manufacturer's protocol. Desalted peptides were dried down again and stored at -20ºC until ready for MS analysis. We chose six proteotypic peptides, listed in **Table S7**, to order as heavy-isotope standards for the absolute quantification of CAT (New England Peptides, Louisville, KY) (7). For LC-MS acquisition, samples were suspended to 500 ng/µL with a 0.1% FA solution containing 5 fmol/µL of each stable-isotope standard peptide. Triplicate injections of 1 µL for each diluted sample were separated on an Acquity µHPLC M-Class Peptide BEH $C_{18}$ column (Waters) using an 80 min segmented linear gradient (Table S8). Peptides were measured on a Q-Exactive HF mass spectrometer (ThermoFisher) using a TOP 10 DDA method with the following parameters: MS1 resolution of 60,000, an automatic gain control (AGC) target of 3e6, and maximum IT of 60 ms; MS2 collected via top 10 DDA with a dynamic exclusion window of 40 s, resolution of 35,000, and AGC target of 1e5.  Raw files were searched using MaxQuant (8) for spectral library generation and peptide quantification was performed with Skyline (9,10). Peptide abundance was determined by the integrated precursor $[M+H]^{2+}$ extracted ion chromatogram and normalized to the heavy isotope standard peptides. Relative CAT abundance was determined as the area response of the SLDDSAYK peptide normalized to the average area response of the RpoB peptides, based on peak shape and coefficient of variance. The WT biological replicate 2 was omitted as an outlier due to mishandling during sample preparation. Enzymatic activity for each construct was determined as the area response of chloramphenicol divided by the relative CAT abundance, then normalized to the mean WT activity.

**Supplementary Figures and Tables**

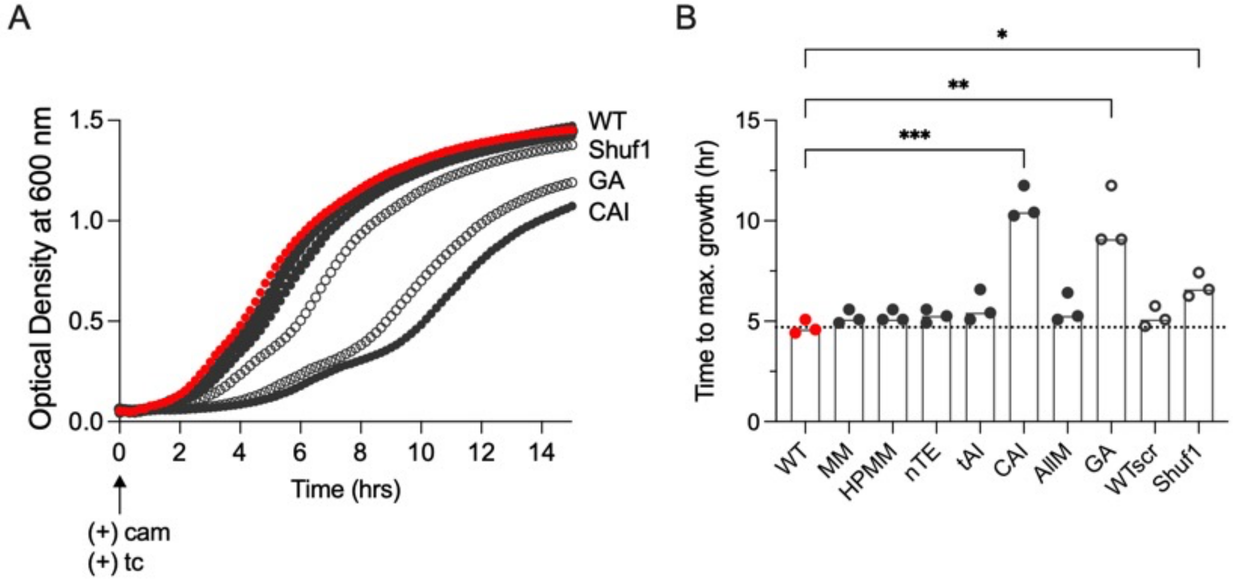


Figure S1. Growth effects on *E. coli* expressing synonymous *cat* sequences. A) Growth of *cat* synonymous sequences induced with 600 ng/mL tetracycline (tc) and challenged with 350 µg/mL chloramphenicol (cam) at the start of growth. B) Quantification of data in panel A as time to maximum growth rate. Growth for GA and CAI synonymous mutants were significantly different from WT; * $p < 0.05$; ** $p < 0.01$; *** $p < 0.001$ (Kruskal-Wallis, uncorrected Dunn’s test).

A

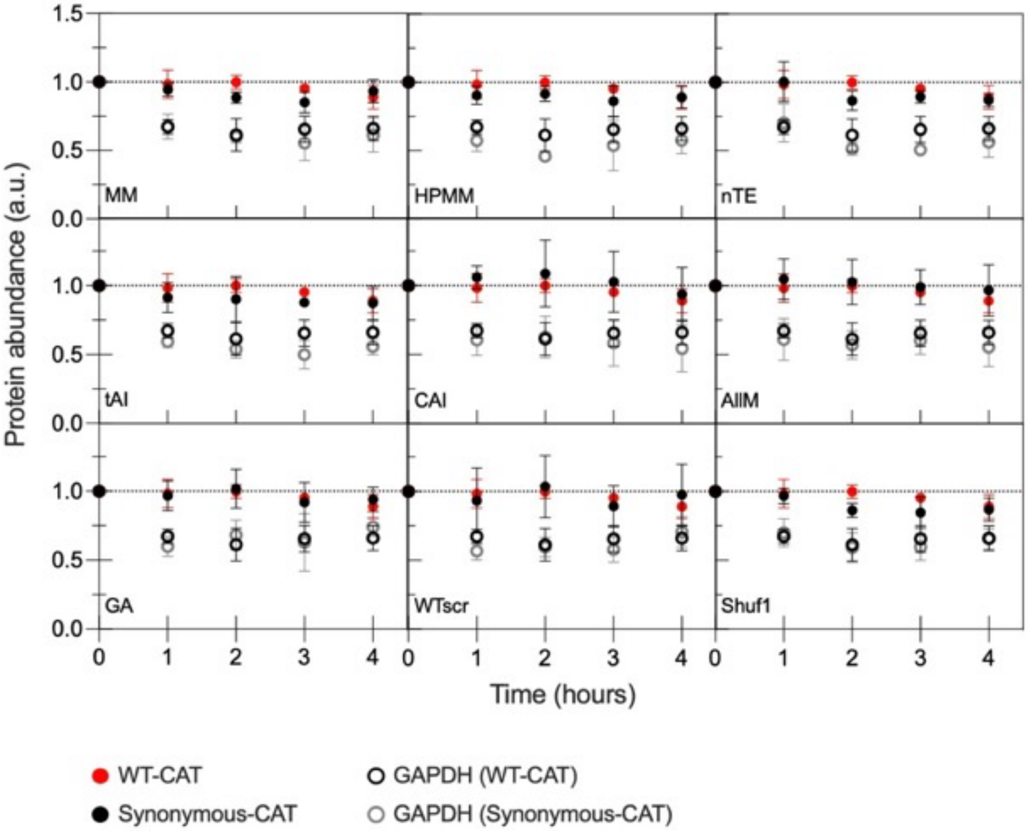


B

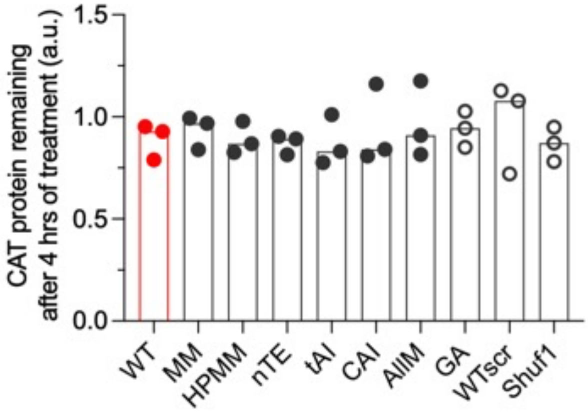


Figure S2. CAT protein turnover in *E. coli*. A) Cells were treated with retapamulin to inhibit CAT protein synthesis and protein was quantified every hour up to 4 hours using fluorescent western blotting. GAPDH was included as a control. B) Amount of CAT protein remaining after 4 hours of retapamulin treatment. Differences were not significant, $p > 0.05$ (Kruskal-Wallis, uncorrected Dunn's test).

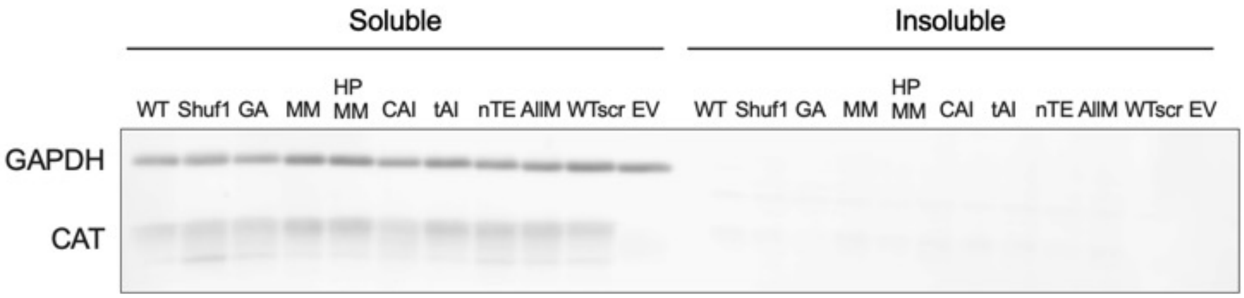


Figure S3. CAT protein solubility and aggregation. To minimize potential variations in solubility due to differences in overall protein accumulation, inducer amount was adjusted to produce similar amounts of CAT protein, as measured by quantitative western blotting. Fluorescent western blot show CAT is present in the soluble but not the insoluble fraction. GAPDH was used as a cell lysis and gel loading control.

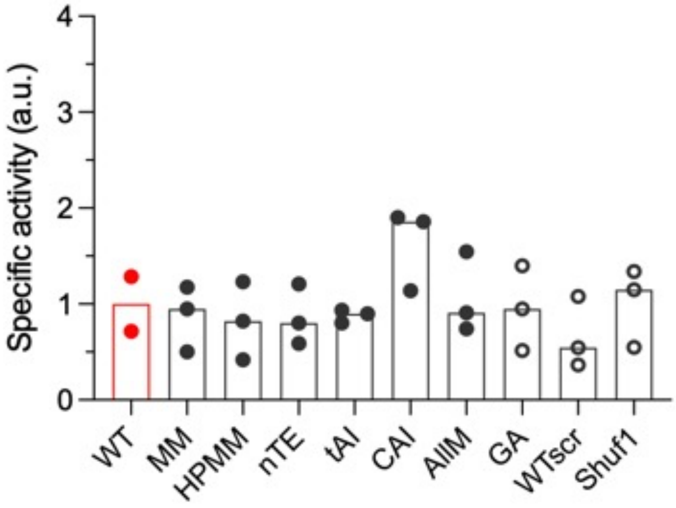


Figure S4. CAT specific activity in whole cell lysates as measured by quantitative mass spectrometry. Specific activity was determined as the area response of CAM divided by CAT abundance, then normalized to WT specific activity (red bar). Bar height indicates the median value of three biological replicates. Closed circles, harmonized synonymous mutants; open circles, non-harmonized synonymous mutants. WT has two biological replicates due to sample mishandling. None of the specific activity measurements were significantly different than WT ($p > 0.05$, Kruskal-Wallis, uncorrected Dunn's test).

Table S1. Codon scores used in CHARMING algorithm

| **Codon** | **MM (usage frequency)** | **High-phi MM (norm 1)** | **CAI** | **tAI** | **nTE** |
|---|---|---|---|---|---|
| TTT | 22.2 | 0.1300 | 0.2960 | 0.1098 | 0.003941 |
| TCT | 8.7 | 1.0000 | 1.0000 | 0.1098 | 0.005343 |
| TAT | 16.5 | 0.2840 | 0.2390 | 0.1646 | 0.006593 |
| TGT | 5.2 | 0.4014 | 0.5000 | 0.0549 | 0.022769 |
| TTC | 15.9 | 1.0000 | 1.0000 | 0.2500 | 0.004732 |
| TCC | 8.9 | 0.7767 | 0.7440 | 0.2500 | 0.011022 |
| TAC | 12.3 | 1.0000 | 1.0000 | 0.3750 | 0.015204 |
| TGC | 6.4 | 1.0000 | 1.0000 | 0.1250 | 0.016196 |
| TTA | 13.8 | 0.0067 | 0.0200 | 0.1250 | 0.008748 |
| TCA | 8.1 | 0.0473 | 0.0770 | 0.1250 | 0.011650 |
| TAA | 2.0 | 1.0000 | 1.0000 | 0.1632 | 1.000000 |
| TGA | 1.1 | 1.0000 | 1.0000 | 0.1250 | 1.000000 |
| TTG | 13.0 | 0.0182 | 0.0200 | 0.1650 | 0.012584 |
| TCG | 8.8 | 0.0897 | 0.0170 | 0.1650 | 0.012573 |
| TAG | 0.3 | 1.0000 | 1.0000 | 0.1632 | 1.000000 |
| TGG | 15.3 | 1.0000 | 1.0000 | 0.1650 | 0.005294 |
| CTT | 11.4 | 0.0263 | 0.0420 | 0.0549 | 0.002855 |
| CCT | 7.2 | 0.0625 | 0.0700 | 0.0549 | 0.006223 |
| CAT | 12.8 | 0.1637 | 0.2910 | 0.0549 | 0.003132 |
| CGT | 20.2 | 1.0000 | 1.0000 | 0.5000 | 0.016996 |
| CTC | 10.5 | 0.0606 | 0.0370 | 0.1250 | 0.007633 |
| CCC | 5.6 | 0.0020 | 0.0120 | 0.1250 | 0.034453 |
| CAC | 9.4 | 1.0000 | 1.0000 | 0.1250 | 0.004332 |
| CGC | 20.8 | 0.3607 | 0.3560 | 0.3600 | 0.013888 |
| CTA | 3.9 | 0.0017 | 0.0070 | 0.1250 | 0.033064 |
| CCA | 8.4 | 0.1297 | 0.1350 | 0.1250 | 0.011501 |
| CAA | 14.7 | 0.1152 | 0.1240 | 0.2500 | 0.018705 |
| CGA | 3.8 | 0.0004 | 0.0040 | 0.0001 | 0.000020 |
| CTG | 51.1 | 1.0000 | 1.0000 | 0.5400 | 0.005768 |
| CCG | 22.4 | 1.0000 | 1.0000 | 0.1650 | 0.004422 |
| CAG | 29.4 | 1.0000 | 1.0000 | 0.3300 | 0.005438 |
| CGG | 6.2 | 0.0003 | 0.0040 | 0.1250 | 0.041105 |
| ATT | 29.7 | 0.2860 | 0.1850 | 0.1646 | 0.003796 |
| ACT | 9.1 | 0.5810 | 0.9650 | 0.1098 | 0.003293 |
| AAT | 19.2 | 0.0556 | 0.0510 | 0.2195 | 0.008135 |
| AGT | 9.4 | 0.0599 | 0.0850 | 0.0549 | 0.008196 |

| Codon | MM (usage frequency) | High-phi MM (norm 1) | CAI | tAI | nTE |
|---|---|---|---|---|---|
| ATC | 23.9 | 1.0000 | 1.0000 | 0.3750 | 0.006851 |
| ACC | 22.8 | 1.0000 | 1.0000 | 0.2500 | 0.004535 |
| AAC | 21.7 | 1.0000 | 1.0000 | 0.5000 | 0.007933 |
| AGC | 16.0 | 1.0000 | 0.4100 | 0.1250 | 0.005525 |
| ATA | 5.5 | 0.0002 | 0.0030 | 0.1632 | 0.040559 |
| ACA | 8.1 | 0.0217 | 0.0760 | 0.1250 | 0.045621 |
| AAA | 34.0 | 1.0000 | 1.0000 | 0.7500 | 0.008856 |
| AGA | 2.9 | 0.0001 | 0.0040 | 0.1250 | 0.027586 |
| ATG | 27.2 | 1.0000 | 1.0000 | 1.0000 | 0.024321 |
| ACG | 15.0 | 0.0983 | 0.0990 | 0.2900 | 0.011957 |
| AAG | 11.0 | 0.2197 | 0.2530 | 0.2400 | 0.012490 |
| AGG | 1.8 | 0.0000 | 0.0020 | 0.1650 | 1.000000 |
| GTT | 18.1 | 1.0000 | 1.0000 | 0.1098 | 0.003697 |
| GCT | 15.4 | 1.0000 | 1.0000 | 0.1098 | 0.005488 |
| GAT | 32.8 | 0.5291 | 0.4340 | 0.1646 | 0.002559 |
| GGT | 24.2 | 1.0000 | 1.0000 | 0.2195 | 0.004942 |
| GTC | 14.8 | 0.2014 | 0.0660 | 0.2500 | 0.014547 |
| GCC | 25.2 | 0.3883 | 0.1220 | 0.2500 | 0.007467 |
| GAC | 19.2 | 1.0000 | 1.0000 | 0.3750 | 0.008251 |
| GGC | 28.1 | 0.7202 | 0.7240 | 0.5000 | 0.008434 |
| GTA | 10.9 | 0.4770 | 0.4950 | 0.6250 | 0.026781 |
| GCA | 20.7 | 0.6825 | 0.5860 | 0.3750 | 0.009936 |
| GAA | 39.3 | 1.0000 | 1.0000 | 0.5000 | 0.007629 |
| GGA | 8.9 | 0.0047 | 0.0100 | 0.1250 | 0.025433 |
| GTG | 26.2 | 0.5095 | 0.2210 | 0.2000 | 0.005151 |
| GCG | 32.3 | 0.8711 | 0.4240 | 0.1200 | 0.001699 |
| GAG | 18.7 | 0.2244 | 0.2590 | 0.1600 | 0.004908 |
| GGG | 11.8 | 0.0263 | 0.0190 | 0.1650 | 0.011141 |

Table S2. Correlation of synonymous *cat* sequences with the WT profile under given codon usage bias (CUB) measure. Correlation of harmonized profiles are in bold.

| **CAT mutant** | **Correlation ($R^2$) with WT profile under given CUB measure** | | | | |
|---|---|---|---|---|---|
| | **%MinMax** | **High-Phi %MinMax** | **CAI** | **tAI** | **nTE** |
| MM | **0.813** | 0.131 | 0.611 | 0.474 | 0.413 |
| HPMM | 0.021 | **0.792** | 0.324 | 0.59 | 0.3 |
| CAI | 0.008 | 0.108 | **0.899** | 0.323 | 0.307 |
| tAI | 0.037 | 0.034 | 0.097 | **0.944** | 0.357 |
| nTE | 0.032 | 0.029 | 0.243 | 0.648 | **0.966** |
| AllM | 0.516 | 0.75 | 0.744 | 0.856 | 0.588 |
| GA | 0.024 | 0.093 | 0.119 | 0.634 | 0.571 |
| WTscr | 0.078 | 0.246 | 0.162 | 0.66 | 0.353 |
| Shuf1 | 0.013 | 0.13 | 0.175 | 0.516 | 0.372 |

Table S3. Primers for FastCloning and site-directed mutagenesis

| Name | Sequence |
|---|---|
| FOR_noTail_allMuts_CAT | TAATCTAGTCAGCTGATCCGGCT |
| REV_WT_noTail | CAGCTGACTAGATTATTTTAATTTACTGTTACACAACTC |
| REV_Shuf1_noTail | CAGCTGACTAGATTACTTCAATTTAGAATTACATAACTCC |
| REV_GA_noTail | CAGCTGACTAGATTATTTCAGTTTGCTGTTACA |
| REV_MMHarm_noTail | CAGCTGACTAGATTATTTCAGCTTAGAATTACATAACTC |
| REV_HPMM_noTail | CAGCTGACTAGATTACTTCAGCTTGGAATTACA |
| REV_CAI_noTail | CAGCTGACTAGATTACTTTAATTTCGAGTTGCAG |
| REV_tAI-Harm_NoTail | CAGCTGACTAGATTATTTCAACTTGGAATTACAAAG |
| REV_nTE_noTail | CAGCTGACTAGATTATTTCAGTTTAGAGTTACACAAC |
| REV_AllModel_noTail | CAGCTGACTAGATTACTTTAACTTAGAATTACATAATTCC |
| REV_scramble_noTail | CAGCTGACTAGATTATTTTAATTTTGAATTACATAATTCTTGT |
| FOR_nfRBS_FC | TGTTCTCTGATATACATATGCATCACCATCACCATCAC |
| REV_nfRBS_FC | GTATATCAGAGAAcaAAAGTTAAACAAAATTATTTCTAGAGGGA AACCG |
| pKT R | ATGTATATCTCCTTCTTAAAGTTAAACAAAATTATTTCTAGAGG GAAA |
| REV_EV pKTS_FC | GAAGGAGATATACATTCTAGTCAGCTGATCCGG |
| FOR_CAT2ndGene_amp | GTTTTTCGCCCTTTGAAAATTGATATCACGACGTTA |
| REV_CAT2ndGene_amp | CAACGTAGCCGGATCAGCTGACTAGATTA |
| FOR_pKTSamp_2ndGene | GATCCGGCTACGTTGGAGTCCACGTTCTTTAATAG |
| REV_pKTSamp_2ndGene | CAACGTAGCCGGATCAGCTGACTAGATTA |

Table S4. gBlocks for FastCloning and MEGAWHOP cloning

| Name | Sequence |
|---|---|
| HPMM-CAT | GTAAAAAATTGGGTTCGCCGTGAGCATTTTGAGTTTTATCGGCATCGT TTACCATGTGGTTTTAGCTTAACAAGCAAAATTGATATCACGACGTTAA AGAAGTCATTGGACGATTCTGCATATAAATTTTATCCTGTAATGATCTA CCTTATAGCGCAGGCAGTAAATCAATTCGACGAGTTACGAATGGCCAT TAAGGACGACGAGTTGATAGTTTGGGACTCCGTTGACCCACAATTCA CGGTTTTCCATCAAGAGACCGAGACCTTTTCTGCTCTCTCATGCCCCT ATAGCTCAGACATAGACCAATTTATGGTCAATTACCTCTCGGTCATGG AGCGCTACAAAAGCGATACTAAGCTTTTTCCACAAGGTGTCACGCCA GAGAATCATTTAAATATAAGCGCATTACCCTGGGTGAATTTCGATTCG TTTAATCTAAATGTAGCCAATTTCACAGATTATTTTGCCCCAATCATCA CCATGGCCAAGTATCAACAAGAAGGAGACCGTTTATTACTTCCGCTGA GTGTGCAGGTTCATCATGCAGTATGTGACGGATTTCACGTAGCGCGA TTCATTAATCGATTACAGGAGCTCTGTAATTCCAAGCTGAAGGCGGCG AACGATGAAAACTATGCGCTGGCGGCGTAATCTAGTCAGCTGATCC |
| CAI-CAT | GTAAAAAATTGGGTTCGCCGTGAGCATTTTGAGTTTTATCGGCATCGT TTACCATGTGGTTTTAGCTTAACAAGCAAAATTGATATCACGACGTTAA AAAAGTCATTAGACGATTCAGCTTACAAATTTTATCCGGTTATGATTTA CTTAATTGCTCAGGCCGTGAACCAATTCGATGAGCTACGAATGGCCAT AAAGGATGACGAGCTGATTGTCTGGGATTCAGTTGATCCGCAGTTTAC AGTGTTCCACCAGGAGACAGAAACATTTTCTGCTCTTAGTTGTCCGTA TAGTAGTGATATTGACCAATTCATGGTTAATTACCTCTCCGTAATGGAA CGGTACAAGTCGGATACGAAACTTTTCCCACAGGGGGTAACGCCTGA GAATCACCTTAACATATCGGCCCTCCCCTGGGTAAACTTCGACTCGTT TAATCTAAATGTAGCAAATTTCACTGATTACTTTGCACCCATCATAACG ATGGCCAAGTATCAACAAGAGGGTGACCGACTCCTTTTACCGCTCTC AGTTCAAGTACATCACGCCGTATGCGACGGTTTCCACGTAGCTCGGT TCATCAATCGACTACAAGAACTCTGCAACTCGAAATTAAAGGCGGCGA ACGATGAAAACTATGCGCTGGCGGCGTAATCTAGTCAGCTGATCC |
| tAI-CAT | GTAAAAAATTGGGTTCGCCGTGAGCATTTTGAGTTTTATCGGCATCGT TTACCATGTGGTTTTAGCTTAACAAGCAAAATTGATATCACGACGTTAA AAAAGTCTCTTGATGACTCTGCCTATAAGTTCTATCCTGTAATGATATA CCTCATAGCACAAGCGGTTAACCAATTCGACGAGCTTAGAATGGCCA TTAAGGATGACGAACTAATTGTATGGGACTCCGTCGACCCTCAGTTCA CTGTATTCCATCAGGAAACCGAAACATTCAGTGCCCTATCCTGTCCAT ATTCGTCGGATATCGACCAGTTTATGGTTAACTATCTGTCAGTTATGG AAAGATATAAAAGTGACACCAAACTTTTCCCGCAGGGTGTGACTCCTG AGAATCACCTCAACATTTCCGCTTTACCATGGGTGAATTTTGATTCTTT TAATCTTAATGTGGCTAATTTTACAGATTATTTCGCCCCGATCATAACT ATGGCAAAGTATCAGCAAGAGGGCGATCGCTTGCTCCTCCCTTTGTC TGTCCAAGTACACCACGCTGTTTGCGACGGATTTCATGTAGCCAGGTT TATAAACAGATTACAGGAGCTTTGTAATTCCAAGTTGAAAGCGGCGAA CGATGAAAACTATGCGCTGGCGGCGTAATCTAGTCAGCTGATCC |

| Name | Sequence |
|---|---|
| AllModel-CAT | GTAAAAAATTGGGTTCGCCGTGAGCATTTTGAGTTTTATCGGCATCGT<br>TTACCATGTGGTTTTAGCTTAACAAGCAAAATTGATATCACGACGTTAA<br>AAAAGTCACTAGATGATTCGGCGTATAAATTCTATCCAGTGATGATTTA<br>TTTAATCGCCCAGGCCGTGAATCAGTTTGACGAGCTCAGGATGGCCA<br>TAAAGGATGATGAACTCATCGTGTGGGACAGTGTTGACCCTCAATTCA<br>CGGTCTTCCATCAGGAGACAGAAACTTTTTCCGCATTGTCGTGTCCAT<br>ACTCCTCAGATATTGATCAGTTTATGGTCAATTATCTCAGCGTTATGGA<br>GCGCTATAAATCAGACACAAAACTATTTCCACAAGGGGTTACTCCCGA<br>GAATCATCTAAATATTTCCGCACTACCGTGGGTGAATTTTGATAGTTTT<br>AATCTAAATGTCGCCAACTTCACAGATTATTTTGCCCCCATAATCACTA<br>TGGCGAAGTACCAACAAGAAGGTGATAGGTTGTTGTTACCACTGAGT<br>GTTCAGGTCCACCACGCTGTGTGTGATGGCTTTCATGTAGCACGGTT<br>TATAAACAGGCTACAGGAATTATGTAATTCTAAGTTAAAGGCGGCGAA<br>CGATGAAAACTATGCGCTGGCGGCGTAATCTAGTCAGCTGATCC |
| GeneArt-CAT | GTAAAAAATTGGGTTCGCCGTGAGCATTTTGAGTTTTATCGGCATCGT<br>TTACCATGTGGTTTTAGCTTAACAAGCAAAATTGATATCACGACGTTAA<br>AAAAGTCATTGGATGATAGCGCCTATAAATTCTATCCGGTGATGATTT<br>ATCTGATTGCCCAGGCAGTTAACCAGTTTGATGAACTGCGTATGGCCA<br>TCAAAGATGATGAGCTGATTGTTTGGGATAGCGTTGATCCGCAGTTTA<br>CCGTTTTTCATCAAGAAACCGAAACCTTTAGCGCACTGAGCTGTCCGT<br>ATAGCAGCGATATTGATCAGTTTATGGTGAACTATCTGAGCGTGATGG<br>AACGCTATAAAAGCGATACCAAACTGTTTCCGCAGGGTGTTACACCG<br>GAAAATCATCTGAATATTTCAGCACTGCCGTGGGTGAACTTTGATTCC<br>TTTAATCTGAATGTGGCCAACTTCACCGATTATTTTGCTCCGATTATTA<br>CCATGGCCAAATATCAGCAAGAAGGTGATCGTCTGCTGCTGCCGCTG<br>AGCGTTCAGGTTCATCATGCAGTTTGTGATGGTTTTCATGTTGCCCGT<br>TTTATCAATCGTCTGCAAGAACTGTGTAACAGCAAACTGAAAGCGGCG<br>AACGATGAAAACTATGCGCTGGCGGCGTAATCTAGTCAGCTGATCC |
| WTscr-CAT | GTAAAAAATTGGGTTCGCCGTGAGCATTTTGAGTTTTATCGGCATCGT<br>TTACCATGTGGTTTTAGCTTAACAAGCAAAATTGATATCACGACGTTAA<br>AAAAGTCATTGGACGATAGCGCATACAAATTCTATCCAGTGATGATTT<br>ATCTAATAGCACAAGCAGTCAACCAGTTCGATGAATTACGTATGGCAA<br>TTAAGGATGATGAGCTCATTGTGTGGGATTCGGTTGATCCGCAGTTTA<br>CAGTTTTTCATCAGGAGACCGAAACCTTCTCCGCGTTATCATGTCCGT<br>ATAGTTCAGATATCGATCAATTTATGGTAAATTATCTGTCAGTTATGGA<br>GCGGTATAAGTCAGATACAAAATTGTTTCCCCAAGGCGTTACCCCTGA<br>AAATCATTTGAATATCAGTGCGTTGCCATGGGTAAATTTTGATTCATTT<br>AATTTGAATGTAGCAAATTTTACAGATTATTTTGCCCCAATAATTACAAT<br>GGCTAAATATCAACAAGAAGGGGATCGCTTACTGCTGCCTTTAAGTGT<br>TCAAGTACATCATGCTGTATGCGATGGGTTTCATGTAGCAAGATTTAT<br>TAATAGATTACAAGAATTATGTAATTCAAAATTAAAAGCGGCGAACGAT<br>GAAAACTATGCGCTGGCGGCGTAATCTAGTCAGCTGATCC |

| Name | Sequence |
| --- | --- |
| nTE-CAT | GTAAAAAATTGGGTTCGCCGTGAGCATTTTGAGTTTTATCGGCATCGT<br>TTACCATGTGGTTTTAGCTTAACAAGCAAAATTGATATCACGACGTTAA<br>AAAAATCCTTAGACGATAGTGCGTATAAGTTTTATCCAGTGATGATCTA<br>CCTTATCGCACAGGCTGTTAACCAATTTGATGAGCTGCGCATGGCCAT<br>AAAAGATGACGAGCTGATCGTATGGGATTCGGTAGACCCTCAATTCA<br>CCGTATTCCACCAAGAAACAGAAACGTTCTCGGCACTCTCGTGTCCG<br>TACTCAAGTGATATCGACCAATTTATGGTGAATTATCTTTCTGTAATGG<br>AGCGGTATAAGTCTGATACCAAGCTATTCCCTCAAGGTGTCACACCTG<br>AAAATCACTTGAATATCAGTGCTCTCCCATGGGTTAACTTTGATTCCTT<br>TAACCTGAACGTTGCTAATTTCACCGACTACTTCGCTCCGATTATAAC<br>AATGGCCAAGTATCAACAAGAGGGTGACCGGTTGCTTTTACCACTTTC<br>CGTGCAAGTGCACCACGCCGTTTGTGATGGTTTTCATGTGGCTCGGT<br>TCATAAATCGGCTCCAGGAGTTGTGTAACTCTAAACTGAAAGCGGCG<br>AACGATGAAAACTATGCGCTGGCGGCGTAATCTAGTCAGCTGATCC |
| MM-CAT | GTAAAAAATTGGGTTCGCCGTGAGCATTTTGAGTTTTATCGGCATCGT<br>TTACCATGTGGTTTTAGCTTAACAAGCAAAATTGATATCACGACGTTAA<br>AAAAGTCATTGGATGATAGCGCTTACAAATTCTACCCGGTCATGATCT<br>ATCTGATTGCTCAGGCCGTGAATCAATTTGACGAACTACGAATGGCGA<br>TAAAAGATGATGAATTAATCGTCTGGGACAGTGTTGATCCTCAATTCA<br>CTGTGTTCCACCAAGAAACTGAGACTTTCAGCGCGTTAAGCTGCCCG<br>TATTCCTCCGATATCGATCAATTTATGGTGAACTACTTAAGCGTGATG<br>GAGCGTTATAAAAGCGATACTAAACTATTCCCCCAAGGAGTTACGCCC<br>GAAAATCATCTAAACATCTCGGCCCTCCCGTGGGTAAACTTTGACAGC<br>TTTAACTTAAACGTCGCTAACTTTACCGATTATTTCGCTCCCATCATCA<br>CAATGGCAAAATACCAACAGGAAGGGGACCGTTTGCTGCTACCGTTG<br>AGCGTACAGGTACATCACGCGGTCTGCGACGGCTTTCATGTGGCCCG<br>TTTCATCAATCGACTCCAAGAGTTATGTAATTCTAAGCTGAAAGCGGC<br>GAACGATGAAAACTATGCGCTGGCGGCGTAATCTAGTCAGCTGATCC |

Table S5. Percent nucleotide changes between synonymous *cat* mutants.

| CAT mutant | WT | Shuf1 | GA | MM | HPMM | CAI | tAI | nTE | AllM | WTscr |
|---|---|---|---|---|---|---|---|---|---|---|
| **WT** | 0.0 | | | | | | | | | |
| **Shuf1** | 18.6 | 0.0 | | | | | | | | |
| **GA** | 16.4 | 19.3 | 0.0 | | | | | | | |
| **MM** | 17.0 | 17.0 | 16.1 | 0.0 | | | | | | |
| **HPMM** | 18.4 | 18.3 | 19.2 | 20.2 | 0.0 | | | | | |
| **CAI** | 19.3 | 18.9 | 19.2 | 20.2 | 17.6 | 0.0 | | | | |
| **tAI** | 20.2 | 17.8 | 19.8 | 20.7 | 18.3 | 20.1 | 0.0 | | | |
| **nTE** | 16.9 | 20.4 | 18.4 | 19.6 | 19.8 | 17.3 | 18.7 | 0.0 | | |
| **AllM** | 17.2 | 17.3 | 17.9 | 17.5 | 18.1 | 20.1 | 17.9 | 19.5 | 0.0 | |
| **WTscr** | 20.5 | 17.5 | 16.7 | 19.8 | 17.5 | 18.7 | 18.6 | 19.6 | 17.8 | 0.0 |

Table S6. Percent codon changes between synonymous *cat* mutants.

| CAT mutant | WT | Shuf1 | GA | MM | HPMM | CAI | tAI | nTE | AllM | WTscr |
|---|---|---|---|---|---|---|---|---|---|---|
| **WT** | 0.0 | | | | | | | | | |
| **Shuf1** | 46.2 | 0.0 | | | | | | | | |
| **GA** | 38.9 | 48.9 | 0.0 | | | | | | | |
| **MM** | 42.5 | 43.0 | 43.9 | 0.0 | | | | | | |
| **HPMM** | 46.6 | 48.9 | 48.4 | 51.1 | 0.0 | | | | | |
| **CAI** | 49.8 | 51.1 | 50.2 | 49.3 | 45.2 | 0.0 | | | | |
| **tAI** | 50.2 | 45.7 | 50.2 | 53.4 | 49.3 | 52.9 | 0.0 | | | |
| **nTE** | 42.5 | 52.5 | 47.1 | 48.9 | 52.5 | 46.2 | 48.0 | 0.0 | | |
| **AllM** | 44.3 | 45.7 | 43.9 | 45.7 | 44.8 | 51.1 | 46.2 | 49.3 | 0.0 | |
| **WTscr** | 49.8 | 47.1 | 39.4 | 48.9 | 47.1 | 47.1 | 47.1 | 51.6 | 43.0 | 0.0 |

Table S7. Peptides used for quantification

| Peptide | Sequence | Light m/z | Heavy m/z | Protein |
|---|---|---|---|---|
| 1 | LPCGFSLTS**K** | 555.2866 | 559.2937 | CAT |
| 2 | SLDDSAY**K** | 449.7113 | 453.7184 | CAT |
| 3 | LQELCNS**K** | 496.2475 | 500.2546 | CAT |
| 3 | AYDLGADV**R** | 490.2458 | 495.2499 | RpoB |
| 4 | ISALGPGGLT**R** | 521.3062 | 526.3103 | RpoB |

Bold sequence letters: isotopically heavy lysine (+8 Da: $^{13}C_6$ $^{15}N_2$) or arginine (+10 Da: $^{13}C_6$ $^{15}N_4$)

Table S8. Liquid chromatography gradient for CAT quantification

| Time (min) | Flow (µL/min) | % ($H_2O$) | % (Acetonitrile + 0.1% formic acid) |
|---|---|---|---|
| Initial | 0.900 | 96.0 | 4.0 |
| 10.00 | 0.900 | 96.0 | 4.0 |
| 61.00 | 0.900 | 70.0 | 30.0 |
| 65.00 | 0.900 | 12.0 | 88.0 |
| 68.00 | 0.900 | 12.0 | 88.0 |
| 69.00 | 0.900 | 96.0 | 4.0 |
| 80.00 | 0.900 | 96.0 | 4.0 |
